# Working memory and reaction time variability mediate the relationship between polygenic risk and ADHD traits in a general population sample

**DOI:** 10.1101/2022.05.31.494251

**Authors:** Mia Moses, Jeggan Tiego, Ditte Demontis, G. Bragi Walters, Hreinn Stefansson, Kari Stefansson, Anders D. Børglum, Aurina Arnatkeviciute, Mark A. Bellgrove

## Abstract

Endophenotypes are heritable and quantifiable traits indexing genetic liability for a disorder. Here, we examined three potential endophenotypes, working memory function, response inhibition and reaction time variability, for attention-deficit hyperactivity disorder (ADHD) measured as a dimensional latent trait in a large general population sample derived from the Adolescent Brain and Cognition Developmental study. The genetic risk for ADHD was estimated using polygenic risk scores (PRS) whereas ADHD traits were quantified as a dimensional continuum using Bartlett factor score estimates, derived from Attention Problems items from the Child Behaviour Checklist and Effortful Control items from the Early Adolescent Temperament Questionnaire-Revised. The three candidate cognitive endophenotypes were quantified using task-based performance measures. Higher ADHD PRSs were associated with higher ADHD traits, as well as poorer working memory performance and increased reaction time variability. Lower working memory performance, poorer response inhibition, and increased reaction time variability were associated with more pronounced ADHD traits. Working memory and reaction time variability partially statistically mediated the relationship between ADHD PRS and ADHD traits, explaining 14% and 16% of the association, respectively. The mediation effect was specific to the genetic risk for ADHD and did not generalise to genetic risk for four other major psychiatric disorders. Together, these findings provide robust evidence from a large general population sample that working memory and reaction time variability can be considered endophenotypes for ADHD that mediate the relationship between ADHD PRS and ADHD traits.

## Introduction

Endophenotypes are heritable quantifiable traits that are argued to index an individual’s genetic liability to develop a given disease or disorder.^1, 2^ Impetus for the identification of endophenotypes for psychiatric disorders was initially driven by earlier failed attempts to identify replicable genetic associations, where the heterogeneity between and within subjects was presumed to swamp the small effects of the genetic signals in relatively small samples. Endophenotypes, such as structural or functional brain imaging or neurocognitive measures, on the other hand, were assumed to have less complex genetic architectures. As a result, it was argued that they should be more closely related to gene function than subjectively rated symptoms of a disorder, and that their use should aid gene discovery.^3, 4^ Although in reality the genetics of the proposed endophenotypes has turned out to be arguably just as complex as the genetics of the disorders themselves,^5–7^ we suggest that the concept of the endophenotype retains utility for understanding the cognitive and neural circuits mediating genetic risk for psychiatric disorders. Here we provide evidence from a large general population cohort – the Adolescent Brain Cognitive Development (ABCD)^8^ study – that cognitive measures of working memory and reaction time variability partially mediate the relationship between polygenic risk for attention deficit hyperactivity disorder (ADHD) and trait measures of attention problems.

Converging evidence to date suggests that the genetic liability for ADHD is driven by both rare and common genetic variation.^9, 10^ Whereas the presence of a single variant in rare cases is sufficient for the development of the disorder, about one-third of the total heritability for ADHD is attributable to common genetic variation that can be quantified through genome wide association studies (GWAS).^9, 11^ Considering this polygenic architecture of ADHD, GWAS discoveries enable us to map the genetic associations between different traits using polygenic risk scores (PRS) that quantify the cumulative genetic risk for a disorder as a weighted sum of disorder-associated single nucleotide polymorphisms (SNPs).^12^ The polygenic risk for ADHD has been associated with a number of specific symptom traits that are linked to ADHD such as hyperactivity, impulsivity, and inattention,^13, 14^ as well as composite ADHD scores.^15, 16^

Cognitive measures that show an association with a disorder, are heritable, and demonstrate evidence of familial overlap, have the potential to serve as endophenotypes for ADHD. Whereas ADHD is a heterogeneous disorder, with any single cognitive mechanism unlikely to be relevant in all cases,^17, 18^ neuropsychological theories consistently highlight the role of executive function impairments associated with ADHD diagnosis.^19–21^ In particular, both children and adults with ADHD tend to demonstrate poorer working memory function,^22, 23^ less efficient response inhibition,^21, 24^ and increased reaction time variability,^25, 26^ the latter likely reflecting a failure of top-down regulation of attention.^27^ Moreover, even in general population samples, individuals with more pronounced ADHD traits tend to experience more difficulties with executive functions.^28^ Twin studies indicate that working memory,^29, 30^ response inhibition,^31, 32^ and reaction time variability,^31, 33^ are moderately heritable with estimates reaching up to h^2^ = 0.7. SNP-heritability studies, quantifying the proportion of phenotypic variance attributable to common genetic variation, also indicate that measures of executive functioning and working memory are significantly heritable,^34, 35^ further confirming the likelihood of additive genetic influences. The potential utility of these cognitive measures as endophenotypes for ADHD is supported by their familial overlap, such that unaffected siblings of individuals with ADHD tend to experience deficits in working memory,^36, 37^ and response inhibition,^38, 39^ and show increased reaction time variability,^38, 39^ as well as broader deficits in executive function.^36^ Support for a genetic overlap between ADHD and cognitive measures have been reported in the recent GWAS meta-analysis of ADHD. In that study the ADHD polygenic risk load (ADHD-PRS) was significantly associated with several measures of cognition, such as degreased attention, working memory and verbal reasoning in 8,722 individuals from the Philadelphia Neurodevelopment Cohort.^11^ Individual differences in executive functions have been found to remain stable across development despite some overall group-level improvements from childhood through to adolescence,^40^ qualifying them as candidate trait-like endophenotypes. Collectively, these results suggest that working memory, response inhibition and reaction time variability may serve as endophenotypes for the dimensional study of ADHD traits.

Mediation models are traditionally used to evaluate the role of potential endophenotypes under the expectation that genetic risk for a disorder operates through the endophenotype.^41^ Although partial mediation, such that some (but not all) of the genetic effects are mediated through the endophenotype provides the most plausible model,^41^ a study in a population sample enriched for ADHD found evidence for full mediation, where working memory and focused attention fully mediated the association between ADHD PRS and hyperactivity-impulsivity, but not inattentive symptoms.^14^ Partial mediation was identified in a sample significantly enriched for cases with ADHD (65% ADHD), where the relationship between ADHD PRS and ADHD status as well as dimensional ADHD symptoms was mediated by working memory and arousal measures.^42^ In a clinical ADHD sample, response inhibition was associated with PRS for ADHD and major depression, and partially mediated the associations with ADHD symptoms,^43^ indicating cross-disorder associations.

Different study designs and sample compositions across studies (population-based enriched for ADHD *vs* clinical *vs* case-control) that all involve a large proportion of subjects with ADHD preclude generalisation of these findings to the broader population. The polygenic architecture of ADHD is thought to explain why it is more recently being considered as the extreme end of a normally distributed continuous trait within the general population.^44, 45^ Therefore, investigating these endophenotypes in relation to dimensional traits of ADHD in a general population would allow one to establish robust associations transcending diagnostic labels. Here, we use a population-based sample from the ABCD^8^ study, which is the largest and most comprehensive longitudinal study of development,^46^ to investigate the potential endophenotypes for ADHD, focusing on working memory function, response inhibition and response time variability. We show that both working memory and response time variability partially mediate the relationship between the genetic risk for ADHD (but not for major depressive disorder or autism spectrum disorder) and its dimensional traits, providing the most robust evidence yet for these measures as endophenotypes for ADHD.

## Methods

### Participants

The present study examined publicly available data from the longitudinal ABCD study (behavioural data – release 3.0, genetic data – release 2.0).^8^ The ABCD study database contains data for up to 11,878 participants aged 9 to 10 years at their baseline assessment. Participants of European ancestry were selected for all further analyses in order to match the genetic ancestry of the discovery genome wide association study (GWAS) for ADHD used to calculate PRSs.^11, 47^ To maximise the sample size for each analysis, three partially overlapping samples at 2-year follow-up were selected to ensure consistency of age across samples (details are provided in the following sections; see Figure 1 for participant demographics). Consistent with the overall prevalence of ADHD in the general population,^48^ 5.3% of all subjects with available parent-rated Child Behaviour Checklist (CBCL)^49^ attention problems scores measured at the 2-year follow-up (*n* = 5,823) would be considered to have clinical ADHD (based on 65 t-score cut-off).^50^

**Figure 1.**
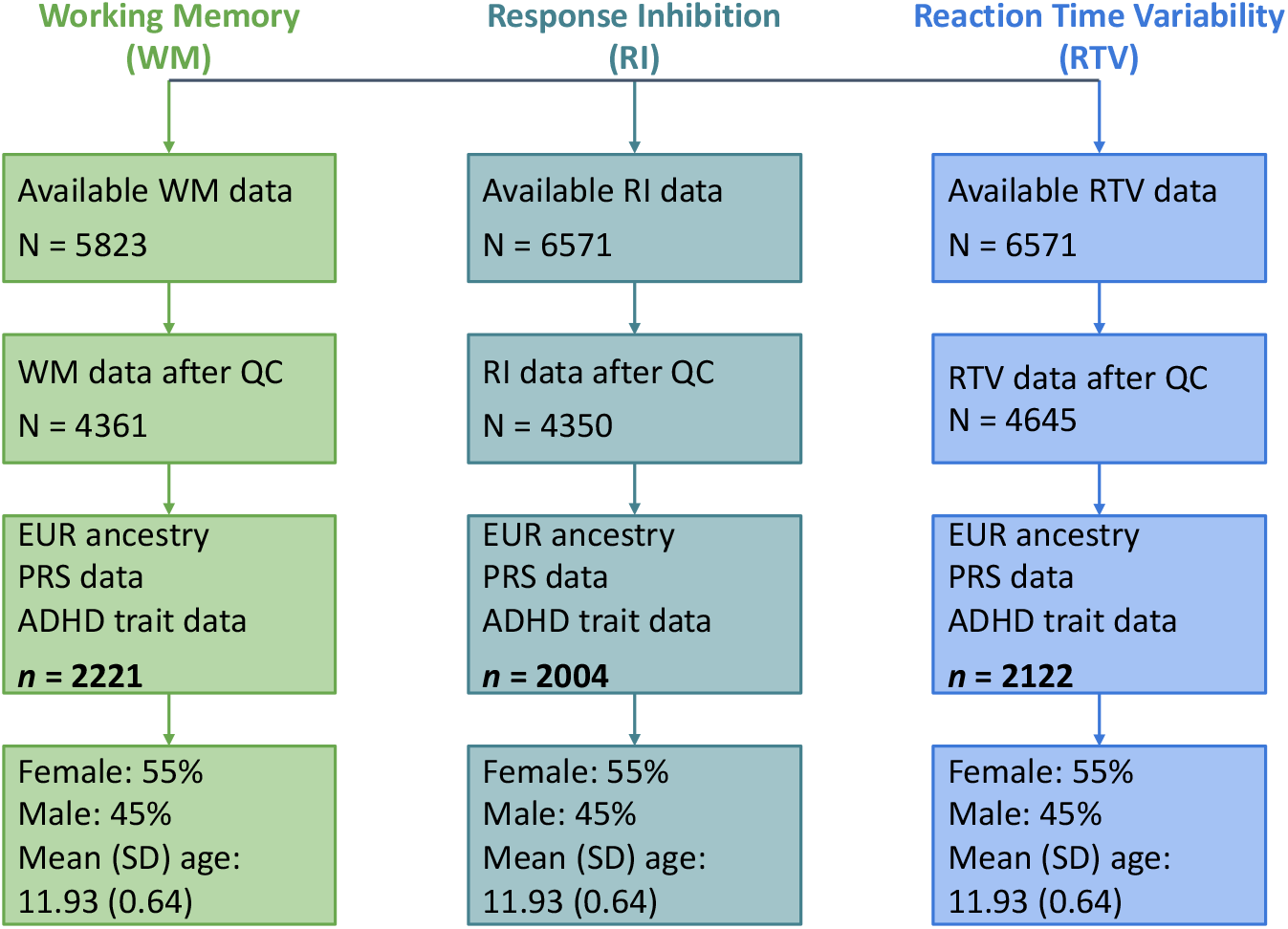
Schematic representation of participant selection process and demographics. Working memory was measured using an emotional N-back task, response inhibition (stop signal reaction time) and reaction time variability were measured with a stop signal task. ADHD traits were quantified as Bartlett factor scores derived from Child Behaviour Checklist Attention Problems items and Early Adolescent Temperament Questionnaire Revised Effortful Control items. Final sample sizes for each measure are in bold. Quality control procedures for each measure are described in section titled “Candidate Cognitive Endophenotypes”. QC = quality control; EUR = European; PRS = polygenic risk score; ADHD = attention-deficit/hyperactivity disorder.

### ADHD Traits Measures

ADHD traits were assessed using the Parent-Rated Child Behaviour Checklist (CBCL)^49^ 6-18 battery and the Early Adolescent Temperament Questionnaire-Revised (EATQ-R)^51^. Both the CBCL and EATQ-R have previously demonstrated clinical utility,^50, 52^ indicating their suitability for quantifying ADHD traits. To increase the accuracy of ADHD trait estimates we performed an Exploratory Factor Analysis (EFA) to extract the factor score estimates that represent more accurate proxies of the true latent scores compared to sum scores (i.e., raw scores).^53^ Factor score estimates are preferable as they address some limitations of raw scores by correcting for error variance and recognising that the strength of the factor loading estimates vary across items.^54, 55^ The 10 Attention Problems items from the CBCL and 18 Effortful Control items from the EATQ-R were included in EFA. Attention Problems items were selected from the CBCL as they have shown prior clinical utility in relation to ADHD.^50^ EATQ-R Effortful Control items were selected from the three-subscale measure as they have been found to be largely unidimensional.^56, 57^ Given phenotypic resolution is often poor at the lower end of clinical scales,^58^ the adaptive end of the CBCL Attention Problems scale was bolstered with the conceptually-related EATQ-R Effortful Control scale to improve phenotypic resolution across the latent trait continuum.^59, 60^ EFA was conducted with a maximum likelihood extraction method and a Promax rotation,^61^ using the ‘psych’ package in RStudio on all participants with available CBCL and EATQ-R data at the 2-year follow-up (*n* = 5,814). Bartlett factor score estimates were created as a continuous measure of ADHD traits, extracting the shared component between measured items, therefore, providing unbiased estimates of the true factor scores, compared to the raw item-based scores.^62^ Lower factor scores indicated more pronounced ADHD traits, and higher factor scores indicated less pronounced ADHD traits. The inverse nature of the scores was attributable to the positive direction of the factor loadings for the EATQ-R Effortful Control items, of which there were a greater number than the CBCL Attention Problems items.

### Polygenic Risk for ADHD

DNA was extracted from saliva samples collected during the baseline visit and genotyped using the Smokescreen Genotyping array for 10,627 subjects.^63, 64^ The following quality control (QC) procedures and genotype imputation were performed prior to ADHD PRS calculation. First, participants with > 10% missing genotype data as well as SNPs with genotyping call rates (GCR) < 90% and SNPs with a minor allele frequency (MAF) of < 0.01 were excluded. Then, several subject-level QC steps were performed by i) stratifying the sample based on ancestry (using self-report and/or based on the first 3 principal components – in which case a sample was assigned the ancestry of the nearest sample from the 1000 Genomes project)^65^; ii) removing subjects with disparities between recorded and observed sex status; iii) removing subjects genotyped on plate number 461 (recommended by the ABCD documentation due to poor data quality); iv) selecting one of the monozygotic twins from the sample by keeping the one with an alphabetically higher subject ID; and v) removing subjects with > 5% missing genotype data, leaving 4988 subjects for imputation. Next, multidimensional scaling was performed using the HapMap3 dataset to identify potential sources of population stratification to be used as covariates in subsequent analyses.^66^ Further, SNPs with GCR <95%, MAF <0.01 and those that significantly departed from the Hardy-Weinberg equilibrium (*p* < 10^-7^) were excluded, leaving 389,183 SNPs for imputation. Imputation was performed using Minimac v4 on the Michigan Imputation Server,^67^ using the reference panel from phase 3 (version 5) of the 1000 Genomes Project Consortium.^65^ After imputation, SNPs with an imputation quality r^2^ > 0.8 and MAF > 0.01 were retained, resulting in a total of 5,958,937 variants.

ADHD PRSs were calculated using the PRSice software package,^68^ based on the GWAS summary statistics for ADHD of 38,691 ADHD cases and 186,843 controls.^11^ The summary statistics were shared by iPSYCH, deCODE and the Psychiatric Genetic Consortium (PGC) based on collaborative grounds prior to their public distribution. Linear regression analyses across a range of p-value thresholds (P_T_) were conducted in PRSice to identify the set of SNPs that maximises the explained variance in ADHD traits (i.e., the ‘optimal’ P_T_)^69^ using age, sex, age^2^, age*sex and three genetic principal components as covariates. PRSice sets a corrected significance level at *p* < .001 to account for multiple comparisons.^68^ PRSs based on the ‘optimal’ P_T_ for each disorder were retained for all future analyses.

### Candidate Cognitive Endophenotype Measures

#### Working memory

Working memory performance in the ABCD study was measured with the emotional N-back task.^8, 70^ The task had low (0-back) and high (2-back) memory load conditions, with task stimuli including three face conditions (positive, neutral, and negative) and one place condition. Participants had to indicate whether a picture presented on a screen was a “Match” or “No Match” on each trial.^8^ In the present study, working memory performance was defined as the mean response accuracy from the two 2-back runs across all four stimulus conditions. All 5,823 ABCD participants had available working memory data at the 2-year follow-up. Prior to inclusion in the analyses, quality control was performed on the working memory data. Participants were included in the present study if i) their overall response accuracy for both 0-back and 2-back conditions was greater than 60% (identified using `tfmri_nback_beh_performflag = 1`); and ii) they had no missing response accuracy scores; leaving 4361 participants. Further, participants of only European ancestry and with available PRS data were selected resulting in a final sample of 2,221 subjects (see Figure 1). The skewness and kurtosis of working memory accuracy scores were −0.70 and −0.13, respectively. An arcsine transformation was applied to working memory accuracy scores to normalise the distribution (skewness = −0.06, kurtosis = −0.32 respectively).

#### Response inhibition

Response inhibition was quantified as the stop signal reaction time (SSRT) from the stop signal task (SST),^8, 71^ derived using the integration method.^72^ The SST required participants to withhold or interrupt a motor response to a “Go” stimulus when it was unpredictably followed by a “Stop” stimulus.^8^ 5823 ABCD participants had available SST data at the 2-year follow-up. Quality control was applied to the SST data such that participants were included in the present study if: i) they had acceptable performance on the task (identified using `tfmri_sst_beh_performflag = 1`); ii) the independent race assumption was not violated such that the mean “Stop Fail” reaction time was greater than mean “Go” reaction time (identified using `tfmri_sst_beh_violatorflag = 0`); iii) no task coding errors occurred (identified using `tfmri_sst_beh_glitchflag = 0`); iv) 25%-75% of all stop trials were performed successfully (identified using `0.25 < tfmri_sst_all_beh_incrs_r < 0.75`); v) “Go” omission rates were less than 30% (identified using `tfmri_sst_all_beh_nrgo_rt < 0.3`); vi) they had stop signal reaction time values higher than 120ms (identified using tfmri_sst_all_beh_total_issrt >= 120); and vii) they had no missing SSRT scores; leaving 4350 participants.^72^ Participants of European ancestry and with available PRS data were selected resulting in a final sample of 2004 subjects (see Figure 1). The skewness and kurtosis of SSRT scores were 0.75 and 1.35, respectively. A log transformation using a natural logarithm was applied to SSRT scores to normalise the distribution (skewness = −0.08, kurtosis = 0.09).

#### Reaction time variability

Reaction time variability (RTV) was quantified as the standard deviation of response times for all correct “Go” trials from the SST.^8, 71^ 5823 ABCD participants had available SST data at the 2-year follow-up. Quality control was applied to the SST data such that participants were included in the present study if i) they had acceptable performance on the task (identified using `tfmri_sst_beh_performflag = 1`); ii) no task coding errors occurred (identified using `tfmri_sst_beh_glitchflag = 0`); iii) “Go” omission rates were less than 30% (identified using `tfmri_sst_all_beh_nrgo_rt < 0.3`); and iv) they had no missing RTV scores; leaving 4645 participants.^72^ Participants of European ancestry with available PRS data were selected resulting a final sample size of 2122 subjects (see Figure 1). The skewness and kurtosis of RTV scores were −0.09 and −0.18, respectively.

### Statistical Analyses

The association between ADHD PRSs and ADHD traits was evaluated through regression analyses in PRSice while controlling for age, sex, age^2^, age*sex as well as three principal components derived based on genetic data as covariates. PRSice sets a corrected significance level at *p* < .001 to account for multiple comparisons.^68^ The associations between each candidate cognitive endophenotype and ADHD traits were evaluated through regression analyses while controlling for age, sex, age^2^ and age*sex as covariates. The associations between ADHD PRSs and each candidate cognitive endophenotype were tested using linear regression while controlling for age, sex, age^2^, age*sex as well as three principal components derived based on genetic data as covariates. In each case, Bonferroni corrections for three regressions (*p* < 0.05/3) were applied to correct for multiple testing.^73^

The possibility that certain cognitive mechanisms lie along the path linking genetic risk for a disorder to symptom traits was examined through mediation analyses assuming that that the genetic risk factor for a psychiatric disorder can operate either fully or partially through the endophenotype.^41^ Mediation analyses were performed if associations between a candidate cognitive endophenotype and both ADHD traits and PRSs were significant (*pcorrected* < 0.05). Statistical mediation was tested using the ‘MeMoBootR’ package in RStudio using age, sex and three genetic principal components as covariates. Bootstrapping method (5,000 bootstrap samples) was used to calculate 95% confidence intervals of indirect effects, where the exclusion of zero indicated a significant indirect effect. To calculate the proportion of the relationship between PRSs and ADHD traits mediated by candidate cognitive endophenotypes, standardized indirect effects were divided by standardized total effects.^74^

Code to reproduce all presented analyses is available in a supplementary file.

## Results

### Exploratory Factor Analysis

Initially, the factorability of the 28 items was examined to justify the application of Exploratory Factor Analysis (EFA).^75^ We estimated that the Kaiser-Meyer-Olkin measure of sampling adequacy (quantifying the proportion of variance among variables that might be caused by underlying factors), was 0.95 – well above the commonly recommended value of 0.50.^76^ Relationships between items were evaluated using Bartlett’s test of sphericity (χ^2^ (378) = 63,437.46,*p* < .001), indicating that correlations between the items were sufficient for EFA.^76^ We then conducted an EFA with maximum likelihood extraction and a Promax (oblique) rotation of the 10 Attention Problems CBCL items and 18 Effortful Control EATQ-R items using data from 5,814 participants. The scree plot exhibited ‘essential unidimensionality’ (see Figure 2 A), as shown by a substantial drop in the eigenvalue between the first and subsequent factors.^77^ Therefore, only one factor was retained (proportion of variance explained: 31.3%) and labelled as ‘ADHD Traits’. Bartlett factor score estimates were created for each participant, quantifying the relative position of each participant along the latent variable continuum of ‘ADHD Traits’, where lower factor scores indicate more pronounced ADHD traits. Bartlett factor score estimates have the highest validity (correlation with underlying factor) compared with other factor score estimates and result in unbiased estimates when analysed with criterion variables.^62^

**Figure 2.**
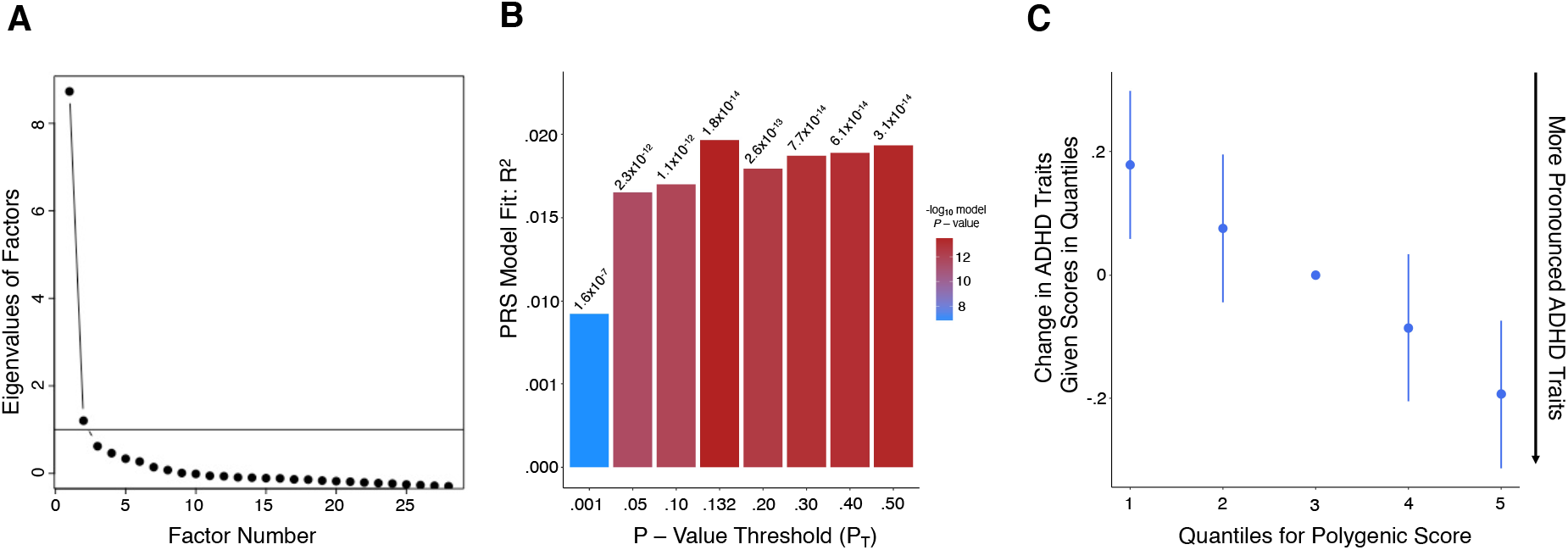
Exploratory factor analysis and the association between ADHD polygenic risk and ADHD traits scores. A) Scree plot from an exploratory factor analysis of CBCL attention problems items and EATQ-R effortful control items indicates ‘essential unidimensionality’. Each dot indicates the amount of common variance of the observed variables each factor explains (i.e., eigenvalue). Any factor with an eigenvalue ?1 explains more variance than a single observed variable (line y = 1), suggesting it should be retained. However, due to the large drop in variance explained by the first factor to the second (and subsequent) factors, a one-factor solution was retained. The total sample used in the EFA, n = 5814. B) Bar plot of the percentage of variance in ADHD trait scores explained by PRSs across a range of p-value thresholds (P_T_), where P_T_ = 0.132 (retaining 51,165 most strongly associated SNPs) explained the most variance in ADHD traits (R^2^ = 2.0%, *p* = 1.8 x 10^-14^). C) Quantile plot demonstrating that as ADHD PRSs increased, ADHD traits increased (i.e., factor scores decreased). The total sample used in the PRS analyses, *n* = 2,847. CBCL = child behaviour checklist; EATQ-R = early adolescence temperament questionnaire revised; ADHD = attention-deficit/hyperactivity disorder; PRS = polygenic risk score.

### Pairwise Associations

First, we evaluated the relationship between ADHD PRSs and ADHD traits. We found that ADHD PRSs were significantly associated with ADHD traits when calculated across a range of P_T_ thresholds with higher ADHD PRSs linked to more pronounced ADHD traits (see Figure 2 A). The highest proportion of variance in ADHD traits was explained at P_T_ = .132 (2.0% variance explained, *p* = 1.8 × 10^-14^), which included 51,165 SNPs associated with ADHD (see Figure 2 B).

We then quantified the associations between ADHD traits and each of the candidate cognitive endophenotypes. Linear regression analyses were adjusted for age, sex, age*sex and age^2^. Reported p-values are corrected for multiple comparisons. Higher working memory function was associated with less pronounced ADHD traits (higher factor scores; *β* [95% *CI*] = 0.18 [0.14, 0.22], *p* = 1.4 × 10^-16^). Lower response inhibition (higher SSRT) was associated with more pronounced ADHD traits (lower factor scores; *β* [95% *CI*] = − 0.13 [− 0.17, − 0.08], *p* = 3.2 × 10^-8^) and increased reaction time variability was associated with more pronounced ADHD traits (lower factor scores; *β* [95% *CI*] = − 0.16 [− 0.20, − 0.12], *p* = 1.2 × 10^-13^). These results suggest that as difficulties in these executive function domains increase, the presence of ADHD behaviours becomes more evident in our population-based sample (Figure 3 A, C, E).

**Figure 3.**
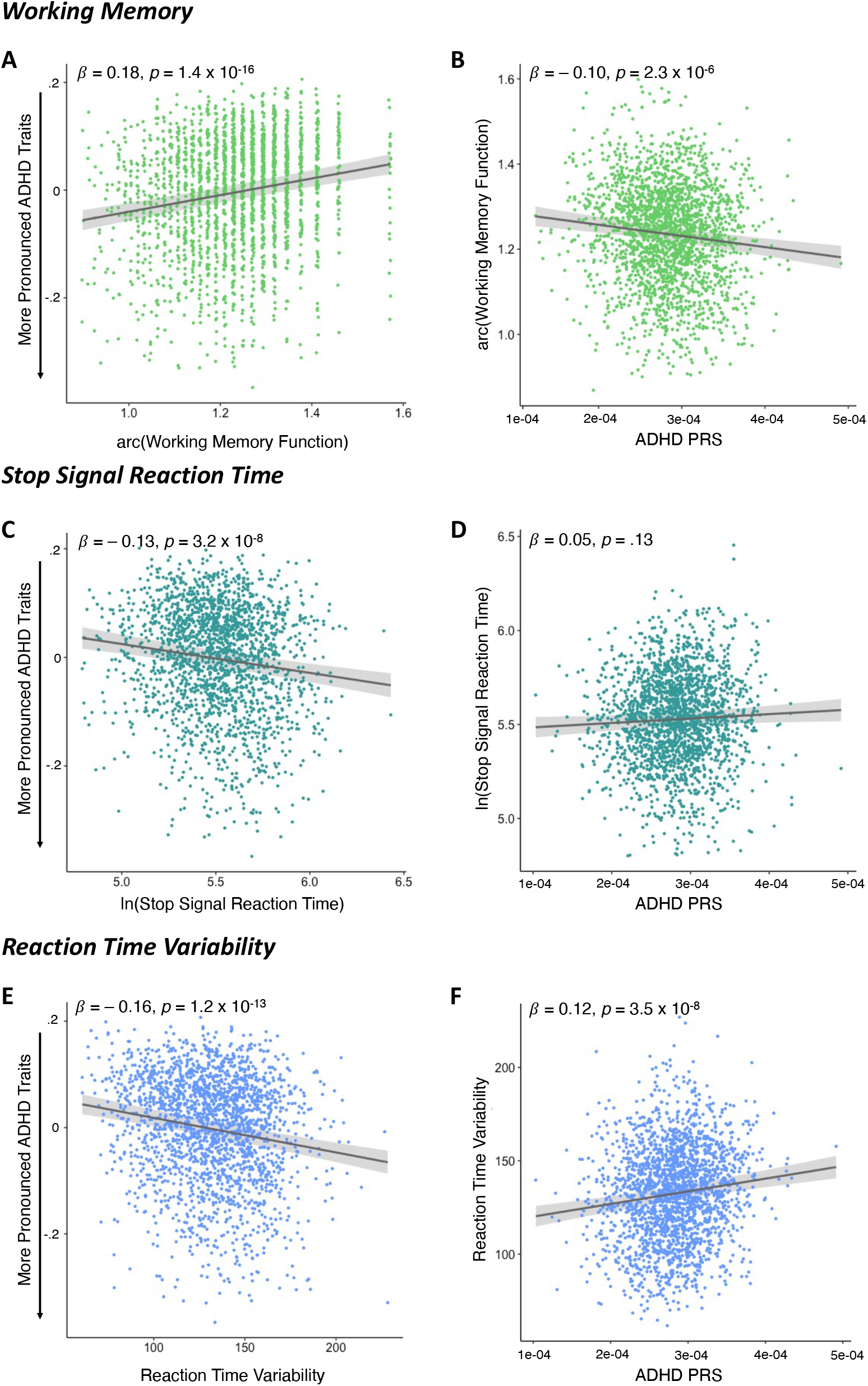
Association between each candidate cognitive endophenotype and ADHD traits (column 1) and between ADHD PRSs and each candidate cognitive endophenotype (column 2). A) Higher working memory accuracy scores form the emotional n-back task were associated with less pronounced ADHD traits (β = 0.18, *p* = 1.4 × 10^-16^, *n* = 2221). B) Higher ADHD PRS were associated with lower working memory response accuracy scores (β = − 0.10, *p* = 2.3 × 10^-6^, *n* = 2221). C) Higher stop signal reaction time scores from the stop signal task were associated with more pronounced ADHD traits (β = − 0.13, *p* = 3.2 × 10^-8^, *n* = 2004). D) No significant association was identified between ADHD PRS and stop signal reaction time scores (β = 0.05, *p* = .13, *n* = 2004). E) Higher reaction time variability scores from the stop signal task were associated with more pronounced ADHD traits (β = − 0.16, *p* = 9.4 × 10^-14^, *n* = 2122). F) Higher ADHD PRS were associated with higher time variability scores (β = 0.12, *p* = 3.4 × 10^-8^, *n* = 2122). ADHD = attention/deficit-hyperactivity disorder; PRS = polygenic risk score. Reported p-values are corrected for multiple comparisons.

To establish that the genetics of ADHD are related to the cognitive measures we evaluated associations between ADHD PRSs (calculated at P_T_ = .132) and candidate cognitive endophenotypes. Linear regression analyses were adjusted for age, sex, age*sex, age^2^ and three genetic principal components to control for the residual effects of population structure. Reported p-values are corrected for multiple comparisons. Higher ADHD PRSs were associated with lower working memory function (*β* [95% *CI*] = − 0.10 [− 0.14, − 0.06], *p* = 2.3 × 10^-6^) and increased reaction time variability (*β* [95% *CI*] = 0.12 [0.08, 0.16],*p* = 3.5 × 10^-8^) (Figure 3 B, F). ADHD PRSs were not associated with response inhibition (*β* [95% *CI*] = 0.05 [0.00, 0.09], *p* = .13) (Figure 3 D). Together, these results indicate that genetic risk for ADHD is associated with executive function difficulties in the working memory and attention/arousal domains, suggesting their potential role as endophenotypes for ADHD.

### Mediation Analyses

To investigate the hypothesis that cognitive traits mediate the relationship between the genetic risk for ADHD and its behavioural manifestations, we tested the statistical mediation for working memory and reaction time variability as these candidate cognitive endophenotypes showed significant associations with both the ADHD traits measure and ADHD PRSs. Mediation analyses were adjusted for age, sex, age*sex, age^2^ and three genetic principal components to control for the residual effects of population structure. We found that both indirect effects via working memory (*b* [95% *CI*] = − 378.70 [− 557.6, − 196.4]) and reaction time variability (*b* [95% *CI*] = − 412.17 [− 594.7, − 235.5]) were significant according to bootstrapping methods. As the direct effects were also statistically significant (Figure 4), both working memory and reaction time variability partially mediated the relationship between the polygenic risk for ADHD and the ADHD trait dimension. The proportion of the association between ADHD PRSs and ADHD traits mediated by working memory performance was 14% (*β*_-0.01716_/ *β*_-0.12476_), whereas reaction time variability mediated 16% (*β*_-0.01859_/ *β*_-0.11977_) of the association. These results provide further evidence for the conceptualisation of working memory and reaction time variability as endophenotypes linking genetic risk for ADHD to its symptom traits.

**Figure 4.**
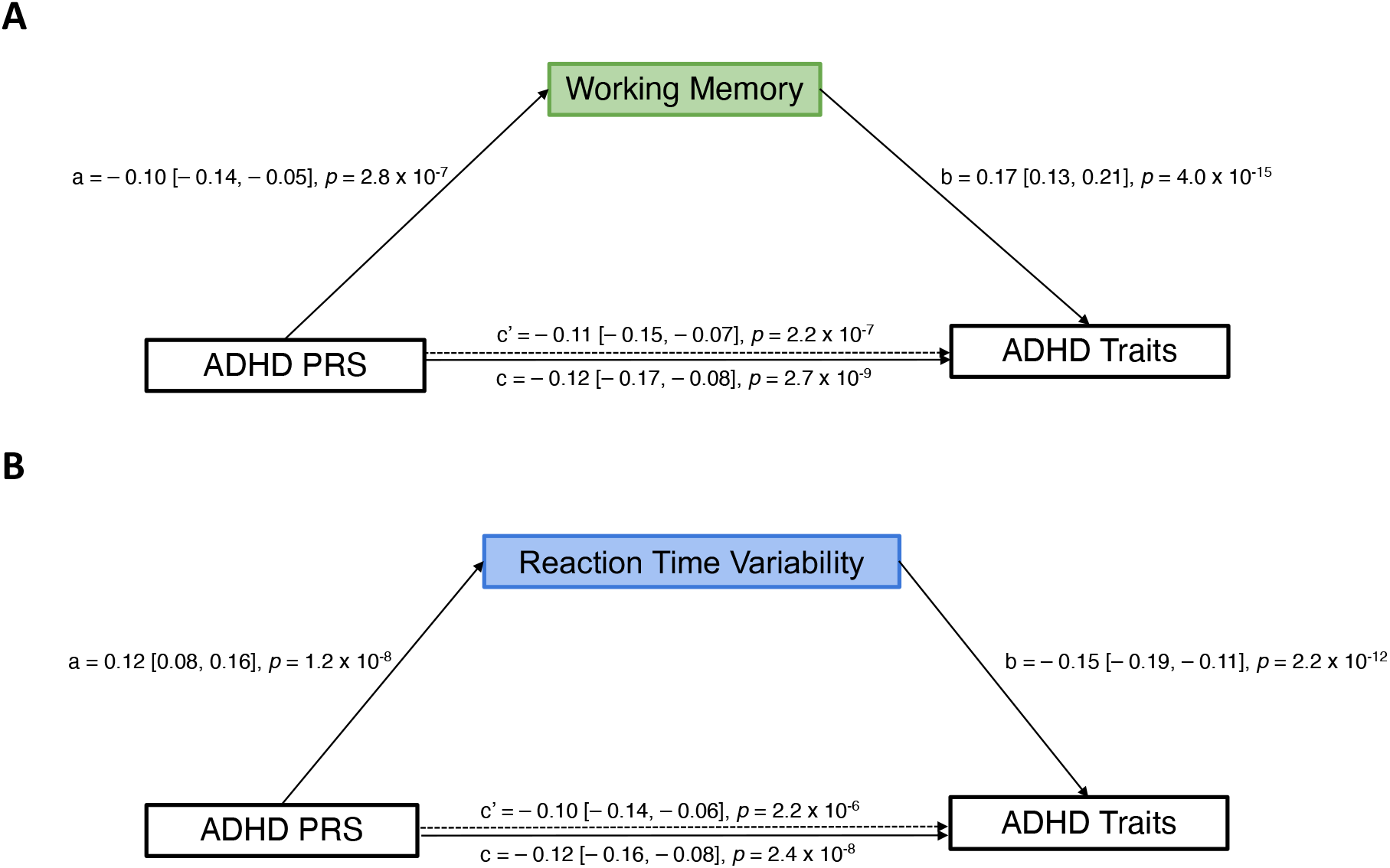
The relationship between ADHD PRS and ADHD traits is partially mediated by working memory and reaction time variability. Standardised regression coefficients and confidence intervals for the relationships between ADHD PRS and ADHD trait factor scores (higher scores = less pronounced ADHD traits) as mediated by candidate cognitive endophenotypes. A) Working memory partially mediated the relationship between ADHD PRS and ADHD traits given both the indirect effect (*b* [95% *CI*] = – 378.70 [– 557.6, – 196.4], *n* = 2221) and the direct effect (i.e., c’ path; β = – 0.11 [– 0.15, – 0.07], *p* = 2.2 x 10^-7^, *n* = 2221) were significant. B) Reaction time variability partially mediated the relationship between ADHD PRS and ADHD traits given both the indirect effect (*b* [95% CI] = – 412.17 [– 594.7, – 235.5], *n* = 2122) and the direct effect (i.e., c’ path; β = – 0.10 [– 0.14, – 0.06], *p* = 2.2 x 10^-6^, *n* = 2122) were significant. ADHD = attention-deficit/hyperactivity disorder; PRS = polygenic risk score. Reported p-values have not been corrected for multiple comparisons.

## Specificity Analyses

The specificity of identified associations was tested by evaluating the associations between PRSs for another four psychiatric disorders [bipolar disorder, schizophrenia, major depressive disorder (MDD) and autism spectrum disorder (ASD)] with the ADHD traits measure and the three cognitive endophenotype measures. First, PRSs for bipolar disorder,^78^ schizophrenia,^79^ MDD,^80^ and ASD,^81^ were created based on GWAS summary statistics downloaded from the PGC (https://www.med.unc.edu/pgc/download-results/). Second, linear regression analyses (using age, sex, age^2^, age*sex and three genetic principal components as covariates) for each disorder across a range of P_T_s were conducted in PRSice to establish whether associations were present with ADHD traits. MDD PRSs at P_T_ = .0945 and ASD PRSs at P_T_ = .003 significantly explained 1.3% (*p* = 8.8 x 10^-10^) and 0.4% (*p* = 5.5 x 10^-4^) of variance in ADHD traits respectively (see Supplementary Figure 1). PRSs for bipolar disorder and schizophrenia did not show significant associations with ADHD traits at any threshold (see Supplementary Figure 1). Next, given their associations with the ADHD traits measure, PRSs for MDD and ASD were tested for their associations with the candidate cognitive endophenotypes using linear regression while controlling for age, sex, age^2^, age*sex as well as three principal components as covariates. Bonferroni correction (*p* < 0.05/6) was applied to account for multiple testing.^73^ No statistically significant associations between MDD PRSs or ASD PRSs and each cognitive measure were found to be significant after corrections for multiple comparisons (*p* < 0.05/6; see Supplementary Figure 2).

## Discussion

Endophenotypes can aid in understanding the cognitive processes mediating genetic risk for psychiatric disorders. ADHD is associated with stable individual differences in executive function that commonly include deficits in working memory, attention, arousal and response inhibition.^19, 20^ Here, we tested a set of candidate cognitive endophenotypes in a large population-based sample and found evidence that working memory and reaction time variability partially mediated the relationship between polygenetic risk for ADHD and a dimensional measure of ADHD traits, supporting their candidacy as endophenotypes. Moreover, the observed mediational relationships were specific to genetic risk for ADHD and not the genetic risk for other major psychiatric disorders.

Considerable evidence links dysfunctional working memory, impaired response inhibition and increased reaction time variability (i.e., attentional control) to ADHD diagnoses.^21, 22, 25^ Our results show that such relationships are also evident in a populationbased sample of children, linking poorer cognitive functioning in each of the three phenotypes with increased ADHD traits. The association between higher ADHD PRSs and poorer working memory accuracy as well as increased reaction time variability are consistent with prior findings where ADHD-PRS was significantly associated with decreased working memory in an independent cohort,^11^ and other studies linking dysfunctions of these mechanisms to ADHD genetic risk factors,^3, 82, 83^ providing further weight to the conceptualisation of working memory and reaction time variability as endophenotypes for ADHD. Partial mediation observed for working memory confirms previous results from a case-control study,^42^ but contrasts the full mediation identified for working memory linking ADHD PRSs and symptoms of hyperactivity/impulsivity,^14^ suggesting that gene-cognition-trait relationships might differ between ADHD symptom domains. Contrary to the well-defined impairments of inhibitory control in ADHD and previous studies indicating associations between genetic risk factors for ADHD and response inhibition,^19, 43, 84^ we found no association between response inhibition and ADHD PRSs in this sample, suggesting alternative genetic pathways may be involved. Measures of self-regulation including SSRT have also been called into question with regards to their utility for investigating interindividual differences,^85^ potentially decreasing the likelihood of identifying associations.

Incorporating data derived from neuroimaging indicating the involvement of relevant structural and/or functional networks into the mediation analyses may provide even further evidence in identifying cognitive endophenotypes. For instance, working memory dysfunction in ADHD is associated with altered front-parietal/striatal network activity.^86, 87^ Therefore, identifying relationships between the genetic risk for ADHD and the neural substrates within these networks would provide additional support for establishing working memory as an endophenotype for ADHD. Serial multiple mediation models may assist in establishing such associations as they investigate the relationship between a predictor and outcome, while modelling the effects of the predictor on the first mediator, which in turn affects the second mediator, thereby affecting the outcome.^88^ Thus far, only one study has found evidence of serial mediation, mapping a pathway from ADHD PRSs to either white matter microstructure of the anterior corona radiata or left dorsomedial prefrontal cortex thickness, then to working memory function, and finally to symptoms of hyperactivity/impulsivity.^14^ Further exploration of these relationships is required to build a fuller and more robust picture of how genetics may be influencing neural substrates, and subsequently the expression of cognitive and behavioural traits associated with ADHD.

The establishment of working memory and reaction time variability as endophenotypes provides clues towards the mechanisms of ADHD psychopathology and can have translational potential, where endophenotypes may eventually act as treatment targets for ADHD through non-pharmacological and pharmacological interventions. Cognitive training to improve working memory function has gained momentum in recent years.^89^ However, when such interventions target only one specific neuropsychological process, there appears to be little clinical value to those with ADHD.^90^ Alongside previously discussed neuropsychological heterogeneity seen in ADHD, ^17, 18^ this suggests the need to develop programs targeting broader ranges of neuropsychological deficits.^90^ The present study therefore suggests that a protocol targeting both working memory and attentional control may accrue greater clinical benefit for individuals with ADHD. Emerging evidence also indicates the possibility to apply transcranial magnetic stimulation, often in conjunction with cognitive training, to areas of the brain associated with neuropsychological deficits to alleviate ADHD symptomology.^91^ Pharmacological treatment for ADHD mainly relies on methylphenidate that acts by blocking the reuptake of dopamine and noradrenaline, increasing their synaptic concentration, and has been shown to improve working memory performance as well as reduce reaction time variability.^92–94^ ADHD has been associated with abnormal processing within attention networks and methylphenidate may act to normalise this.^95^ Moreover, methylphenidate-induced improvements in working memory performance occur with task-related reductions in regional cerebral blood flow in the dorsolateral prefrontal cortex and posterior parietal cortex.^96^ Pivotal work in non-human primates suggests that working memory related impairments in ADHD likely manifest through disruption to α2 adrenoceptor and the dopamine D1 receptors signalling in prefrontal cortical areas,^97^ thus providing a mechanistic bases for cognitive deficits in ADHD.

Our results should be considered in light of some limitations. Here we evaluated the genetic risk for ADHD based on case-control GWAS and used it to assess dimensional ADHD traits in a population-based sample, which might weaken the identified associations.^98, 99^ GWASs on large population-based samples for ADHD traits,^100^ may therefore offer an additional direction for future research in this field. Additionally, expanding the scope of the genetic research by incorporating data from diverse ancestries will improve the generalisability of these results and provide better representation of population-based samples.

Together, our results provide compelling evidence that working memory and reaction time variability partially statistically mediate the relationship between the genetic risk for ADHD and its dimensional traits in a large population-based sample. These findings support the conceptualisation of working memory and reaction time variability as endophenotypes for ADHD and offer a mechanistic basis on which both pharmacological and non-pharmacological interventions may be targeted to reduce the influence of genetic liability on ADHD symptomatology.

## Supporting information

Code to reproduce results

Code and plots in html format

## Acknowledgements

MAB was supported by a National Health and Medical Research Council of Australia Senior Research Fellowship. AA was funded by a grant from the Australian Research Council (ARC) under its Linkage Project scheme (LP160101592). JT was supported by a Turner Impact Fellowships from the Turner Institute for Brain and Mental Health at Monash University. Data used in the preparation of this article were obtained from the Adolescent Brain Cognitive Development^SM^ (ABCD) Study (https://abcdstudy.org), held in the NIMH Data Archive (NDA). This is a multisite, longitudinal study designed to recruit more than 10,000 children age 9-10 and follow them over 10 years into early adulthood. The ABCD Study^®^ is supported by the National Institutes of Health and additional federal partners under award numbers U01DA041048, U01DA050989, U01DA051016, U01DA041022, U01DA051018, U01DA051037, U01DA050987, U01DA041174, U01DA041106, U01DA041117, U01DA041028, U01DA041134, U01DA050988, U01DA051039, U01DA041156, U01DA041025, U01DA041120, U01DA051038, U01DA041148, U01DA041093, U01DA041089, U24DA041123, U24DA041147. A full list of supporters is available at https://abcdstudy.org/federal-partners.html. A listing of participating sites and a complete listing of the study investigators can be found at https://abcdstudy.org/consortium_members/. ABCD consortium investigators designed and implemented the study and/or provided data but did not necessarily participate in the analysis or writing of this report. This manuscript reflects the views of the authors and may not reflect the opinions or views of the NIH or ABCD consortium investigators. The ABCD data repository grows and changes over time. The ABCD genetic data used in this report came from ABCD release 2.0; https://nda.nih.gov/study.html?id=634. The ABCD behavioural/cognitive data used in this report came from ABCD release 3.0; https://nda.nih.gov/study.html?id=901.

