## Supplementary figures for "Working memory and reaction time variability mediate the relationship between polygenic risk and ADHD traits in a general population sample"

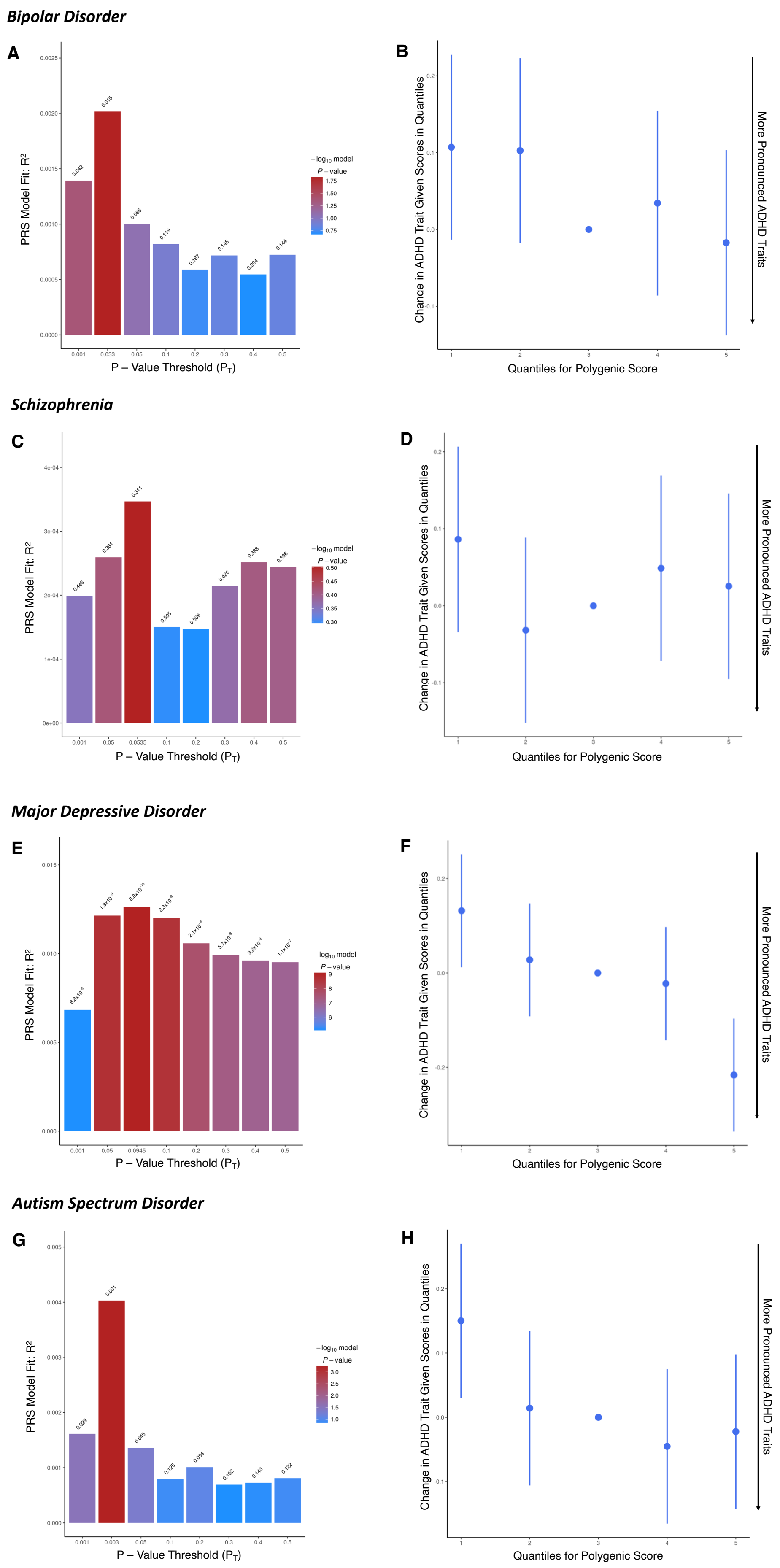

Bipolar Disorder
A
PRS Model Fit: R²
P – Value Threshold (P_T)
−log10 model P – value
B
Change in ADHD Trait Given Scores in Quantiles
Quantiles for Polygenic Score
More Pronounced ADHD Traits
Schizophrenia
C
PRS Model Fit: R²
P – Value Threshold (P_T)
−log10 model P – value
D
Change in ADHD Trait Given Scores in Quantiles
Quantiles for Polygenic Score
More Pronounced ADHD Traits
Major Depressive Disorder
E
PRS Model Fit: R²
P – Value Threshold (P_T)
−log10 model P – value
F
Change in ADHD Trait Given Scores in Quantiles
Quantiles for Polygenic Score
More Pronounced ADHD Traits
Autism Spectrum Disorder
G
PRS Model Fit: R²
P – Value Threshold (P_T)
−log10 model P – value
H
Change in ADHD Trait Given Scores in Quantiles
Quantiles for Polygenic Score
More Pronounced ADHD Traits

Supplementary Figure 1 **Associations between polygenic risk scores for bipolar disorder (A,B), schizophrenia (C,D), major depressive disorder (E,F), autism spectrum disorder (G,H) and ADHD trait scores.** First column in each subplot – bar plot showing the associations between ADHD trait scores and PRS across a range of p-value thresholds ($P_T$). Second column – quantile plot demonstrating the direction of the identified association at the best-fitting $P_T$. No significant associations were identified for bipolar disorder (A,B) and schizophrenia (C,D) PRS. PRS for major depressive disorder ($p_T$ = 0.0945, R2 = 1.3%, p = 8.8 x $10^{-10}$) and autism spectrum disorder ($p_T$ = 0.003, R2 = 0.4%, p = 0.001) was associated with ADHD traits, where increased PRS was associated with increased ADHD traits (decreased factor scores). The total sample used in the PRS analyses, n = 2847. PRS = polygenic risk score.

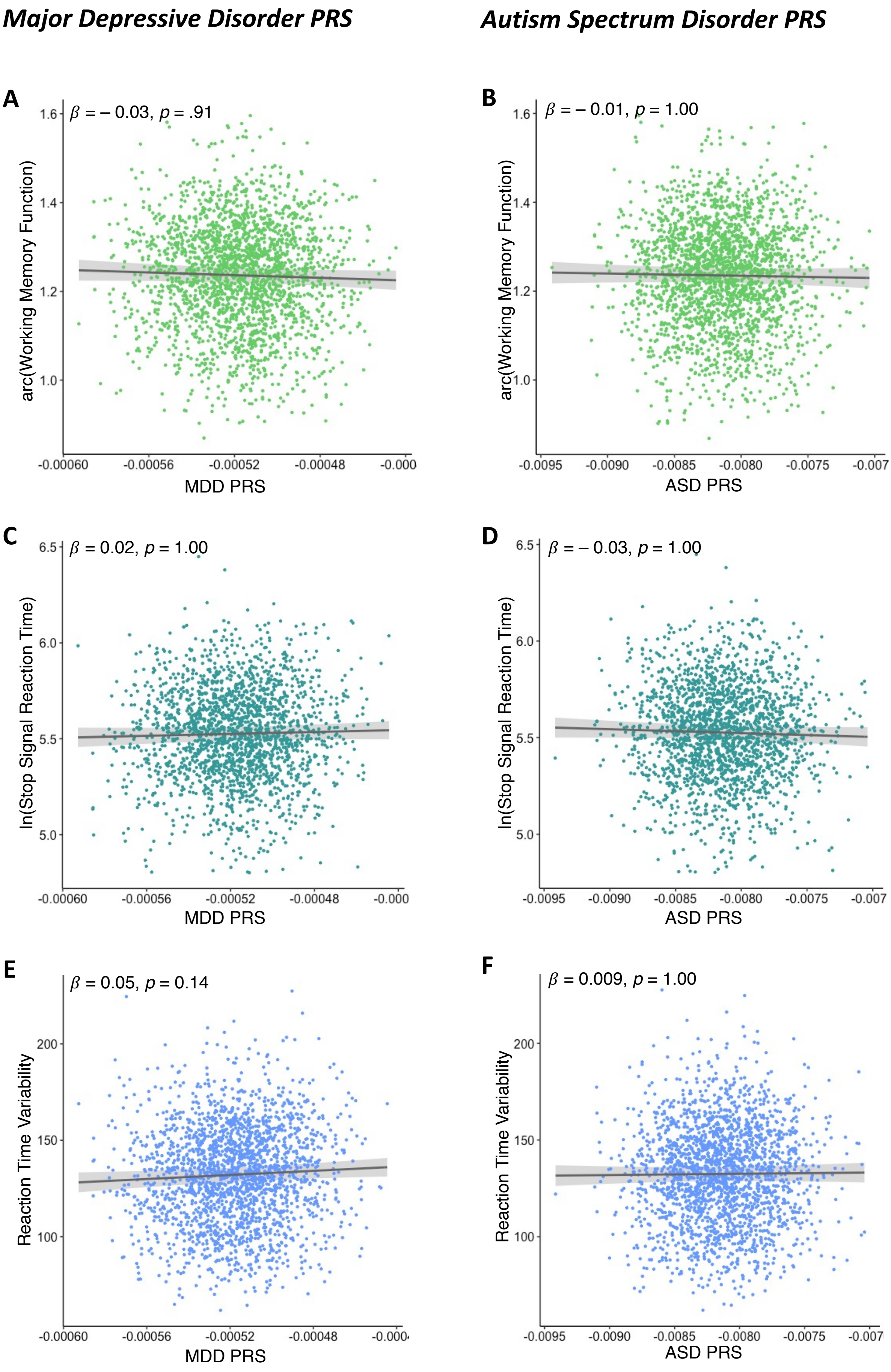

Major Depressive Disorder PRS
Autism Spectrum Disorder PRS
A
β = – 0.03, p = .91
arc(Working Memory Function)
MDD PRS
B
β = – 0.01, p = 1.00
arc(Working Memory Function)
ASD PRS
C
β = 0.02, p = 1.00
ln(Stop Signal Reaction Time)
MDD PRS
D
β = – 0.03, p = 1.00
ln(Stop Signal Reaction Time)
ASD PRS
E
β = 0.05, p = 0.14
Reaction Time Variability
MDD PRS
F
β = 0.009, p = 1.00
Reaction Time Variability
ASD PRS

Supplementary Figure 2 **Associations between three ADHD candidate cognitive endophenotypes and MDD PRSs (column 1) and ASD PRSs (column 2).** No significant association was identified between MDD PRS and working memory accuracy scores form the emotional n-back task ($\beta = -0.03$, $p = 0.91$, $n = 2221$, A); stop signal reaction time scores from the stop signal task ($\beta = 0.02$, $p = 1.00$, $n = 2004$, C); reaction time variability scores from the stop signal task ($\beta = 0.05$, $p = 0.14$, $n = 2122$, E) as well as between ASD PRS and PRS working memory accuracy scores ($\beta = -0.01$, $p = 1.00$, $n = 2221$, B); stop signal reaction time scores ($\beta = -0.03$, $p = 1.00$, $n = 2004$, D) and reaction time variability scores ($\beta = 0.009$, $p = 1.00$, $n = 2122$, F). All p-values are Bonferroni corrected for multiple comparisons. PRS = polygenic risk score; MDD = major depressive disorder; ASD = autism spectrum disorder.
