## Supplementary material for "Working memory and reaction time variability mediate the relationship between polygenic risk and ADHD traits in a general population sample": Code and plots in html format: Moses_Endophenotypes_HTMLcode_results_20220601.html

### Load packages

```
library(data.table)
library(ggplot2)
library(ggpubr)
library(visreg)
library(mice)
```

```
## 
## Attaching package: 'mice'
```

```
## The following object is masked from 'package:stats':
## 
##     filter
```

```
## The following objects are masked from 'package:base':
## 
##     cbind, rbind
```

```
library(lme4)
```

```
## Loading required package: Matrix
```

```
## Registered S3 methods overwritten by 'lme4':
##   method                          from
##   cooks.distance.influence.merMod car 
##   influence.merMod                car 
##   dfbeta.influence.merMod         car 
##   dfbetas.influence.merMod        car
```

```
library(lmerTest)
```

```
## 
## Attaching package: 'lmerTest'
```

```
## The following object is masked from 'package:lme4':
## 
##     lmer
```

```
## The following object is masked from 'package:stats':
## 
##     step
```

```
library(JWileymisc)
library(multilevelTools)
library(emmeans)
library(MeMoBootR)
```

```
## Loading required package: boot
```

```
## Loading required package: diagram
```

```
## Loading required package: shape
```

```
library(FSA)
```

```
## Registered S3 methods overwritten by 'FSA':
##   method       from
##   confint.boot car 
##   hist.boot    car
```

```
## ## FSA v0.9.1. See citation('FSA') if used in publication.
## ## Run fishR() for related website and fishR('IFAR') for related book.
```

```
library(lm.beta)
library(sjmisc)
library(pscl)
```

```
## Classes and Methods for R developed in the
## Political Science Computational Laboratory
## Department of Political Science
## Stanford University
## Simon Jackman
## hurdle and zeroinfl functions by Achim Zeileis
```

```
library(maczic)
```

```
## Loading required package: MASS
```

```
library(mediation)
```

```
## Loading required package: mvtnorm
```

```
## Loading required package: sandwich
```

```
## mediation: Causal Mediation Analysis
## Version: 4.5.0
```

```
library(foreign)
library(MASS)
library(AER)
```

```
## Loading required package: car
```

```
## Loading required package: carData
```

```
## 
## Attaching package: 'car'
```

```
## The following object is masked from 'package:FSA':
## 
##     bootCase
```

```
## The following object is masked from 'package:boot':
## 
##     logit
```

```
## Loading required package: lmtest
```

```
## Loading required package: zoo
```

```
## 
## Attaching package: 'zoo'
```

```
## The following objects are masked from 'package:base':
## 
##     as.Date, as.Date.numeric
```

```
## Loading required package: survival
```

```
## 
## Attaching package: 'survival'
```

```
## The following object is masked from 'package:boot':
## 
##     aml
```

```
library(boot)
library(performance)
library(MBESS)
```

```
## 
## Attaching package: 'MBESS'
```

```
## The following objects are masked from 'package:JWileymisc':
## 
##     cor2cov, smd
```

```
library(EnvStats)
```

```
## 
## Attaching package: 'EnvStats'
```

```
## The following object is masked from 'package:MBESS':
## 
##     cv
```

```
## The following object is masked from 'package:car':
## 
##     qqPlot
```

```
## The following object is masked from 'package:MASS':
## 
##     boxcox
```

```
## The following object is masked from 'package:Matrix':
## 
##     print
```

```
## The following objects are masked from 'package:stats':
## 
##     predict, predict.lm
```

```
## The following object is masked from 'package:base':
## 
##     print.default
```

```
library(EFA.dimensions)
library(psych)
```

```
## 
## Attaching package: 'psych'
```

```
## The following object is masked from 'package:MBESS':
## 
##     cor2cov
```

```
## The following object is masked from 'package:car':
## 
##     logit
```

```
## The following object is masked from 'package:mediation':
## 
##     mediate
```

```
## The following object is masked from 'package:FSA':
## 
##     headtail
```

```
## The following object is masked from 'package:boot':
## 
##     logit
```

```
## The following object is masked from 'package:JWileymisc':
## 
##     cor2cov
```

```
## The following objects are masked from 'package:ggplot2':
## 
##     %+%, alpha
```

```
library(plyr)
```

```
## 
## Attaching package: 'plyr'
```

```
## The following object is masked from 'package:FSA':
## 
##     mapvalues
```

```
## The following object is masked from 'package:ggpubr':
## 
##     mutate
```

### Load data

```
# Polygenic risk scores (PRS) for ADHD
data_PRS_ADHD <- read.table("ADHD_PRSice_SCORES_AT_ALL_THRESHOLDS.txt",
                       header = TRUE, sep = " ", stringsAsFactors = FALSE) 
names(data_PRS_ADHD)[1] <- 'src_subject_id' 

# PRS for Autism Spectrum Disorder
data_PRS_AUT <- read.table("AUT_PRSice_SCORES_AT_ALL_THRESHOLDS.txt",
                       header = TRUE, sep = " ", stringsAsFactors = FALSE) 
names(data_PRS_AUT)[1] <- 'src_subject_id' 

# PRS for Bipolar Disorder
data_PRS_BIP <- read.table("BIP_PRSice_SCORES_AT_ALL_THRESHOLDS.txt",
                       header = TRUE, sep = " ", stringsAsFactors = FALSE) 
names(data_PRS_BIP)[1] <- 'src_subject_id' 

# PRS for Major Depressive Disorder
data_PRS_MDD <- read.table("MDD_PRSice_SCORES_AT_ALL_THRESHOLDS.txt",
                       header = TRUE, sep = " ", stringsAsFactors = FALSE) 
names(data_PRS_MDD)[1] <- 'src_subject_id' 

# PRS for Schizophrenia
data_PRS_SCZ <- read.table("SCZ_PRSice_SCORES_AT_ALL_THRESHOLDS.txt",
                       header = TRUE, sep = " ", stringsAsFactors = FALSE) 
names(data_PRS_SCZ)[1] <- 'src_subject_id' 

# Genetic PCs covariates (PC1, PC2, PC3)
data_PCs <- read.table("ABCD_eur_covariatesPC3_PLINK.txt",
                       header = TRUE, sep = "\t", stringsAsFactors = FALSE)
names(data_PCs)[1] <- 'src_subject_id' 
data_PCs <- dplyr::select(data_PCs, c('src_subject_id','PC1', 'PC2', 'PC3'))

# Child Behaviour Checklist (CBCL)
data_CBCL <- read.table("abcd_cbcls01.txt", header = TRUE, sep = "\t", 
                    stringsAsFactors = FALSE)

# Early Adolescent Teperament Questionnarie Revised (EATQ-R) and CBCL items
data_EATQR_CBCL <- read.table("CBCL_EATQ_BISBAS_2year.txt", header = TRUE, 
                              sep = "\t", stringsAsFactors = FALSE)

# Working Memory (WM)
data_WM <- read.table("abcd_mrinback02.txt",
                       header = TRUE, sep = "\t", stringsAsFactors = FALSE)

# Stop Signal Reaction Time (SSRT)
data_SSRT <- read.table("abcd_sst02.txt",
                       header = TRUE, sep = "\t", stringsAsFactors = FALSE)

# Reaction Time Variability (RTV)
data_RTV <- read.table("abcd_sst02.txt",
                       header = TRUE, sep = "\t", stringsAsFactors = FALSE)
```

### Select 2-year follow-up and change character to numeric

```
# CBCL at 2-year follow-up
data_CBCL_2y <- filter(data_CBCL, eventname == "2_year_follow_up_y_arm_1")
data_CBCL_2y[, 10:79] <- lapply(data_CBCL_2y[, 10:79], as.numeric)

# WM at 2-year follow-up 
data_WM_2y <- filter(data_WM, eventname == "2_year_follow_up_y_arm_1")
data_WM_2y[, 13:590] <- lapply(data_WM_2y[, 13:590], as.numeric)

# SSRT at 2-year follow-up 
data_SSRT_2y <- filter(data_SSRT, eventname == "2_year_follow_up_y_arm_1")
data_SSRT_2y[, 13:104] <- lapply(data_SSRT_2y[, 13:104], as.numeric)

# RTV at 2-year follow-up
data_RTV_2y <- filter(data_RTV, eventname == "2_year_follow_up_y_arm_1")
data_RTV_2y[, 13:104] <- lapply(data_RTV_2y[, 13:104], as.numeric)
```

### Quality control measures

```
# Quality control for WM data 
data_WM_2y_QC <- filter(data_WM_2y, 
                        tfmri_nback_beh_performflag == 1, # performance good
                        !is.na(tfmri_nb_all_beh_c2b_rate)) # remove missing

# Quality control for SSRT data
data_SSRT_2y_QC <- filter(data_SSRT_2y, 
                          tfmri_sst_beh_performflag == 1, # performance good
                          tfmri_sst_beh_violatorflag == 0, # racing assum met
                          tfmri_sst_beh_glitchflag == 0, # no issues run task
                          between(tfmri_sst_all_beh_incrs_rt, .25, .75), # prop incor 
                          tfmri_sst_all_beh_nrgo_rt < .3, # go omission < 30%
                          tfmri_sst_all_beh_total_issrt>=120, # reliable scores
                          !is.na(tfmri_sst_all_beh_total_issrt)) # remove missing

# Quality control for RTV data
data_RTV_2y_QC <- filter(data_RTV_2y, 
                          tfmri_sst_beh_performflag == 1, # performance good
                          tfmri_sst_beh_glitchflag == 0, # no issues run task
                          tfmri_sst_all_beh_nrgo_rt < .3, # go omission < 30%
                          !is.na(tfmri_sst_all_beh_crgo_stdrt)) # remove missing
```

### Run exploratory factor analysis to create an outcome variable

```
# Data selection - Effortful control (EC) items from the EATQ-R and AP items from CBCL
# note - this data is already at the 2 year follow-up
data_EFA <- dplyr::select(data_EATQR_CBCL, c('src_subject_id','interview_age', 'sex', 'cbcl_q01_p', 'cbcl_q04_p', 'cbcl_q08_p', 'cbcl_q10_p', 'cbcl_q13_p', 'cbcl_q17_p', 'cbcl_q41_p', 'cbcl_q61_p', 'cbcl_q78_p', 'cbcl_q80_p', 'eatq_finish_p', 'eatq_deal_p', 'eatq_turn_taking_p', 'eatq_open_present_p', 'eatq_before_hw_p', 'eatq_concentrate_p', 'eatq_right_away_p', 'eatq_distracted_p', 'eatq_impulse_p', 'eatq_try_focus_p', 'eatq_finish_hw_p', 'eatq_early_start_p', 'eatq_peripheral_p', 'eatq_puts_off_p', 'eatq_laugh_control_p', 'eatq_sidetracked_p','eatq_stick_to_plan_p', 'eatq_close_attention_p'))

# remove missing data
comp_data_EFA <- data_EFA[complete.cases(data_EFA), ]
# take out subject ID, age and sex
data_for_EFA <- comp_data_EFA[, c(4:31)]

# Kaiser-Meyer-Olkin test
KMO(data_for_EFA) # we have enough participants
```

```
## Kaiser-Meyer-Olkin factor adequacy
## Call: KMO(r = data_for_EFA)
## Overall MSA =  0.95
## MSA for each item = 
##             cbcl_q01_p             cbcl_q04_p             cbcl_q08_p 
##                   0.98                   0.97                   0.94 
##             cbcl_q10_p             cbcl_q13_p             cbcl_q17_p 
##                   0.94                   0.91                   0.94 
##             cbcl_q41_p             cbcl_q61_p             cbcl_q78_p 
##                   0.95                   0.97                   0.95 
##             cbcl_q80_p          eatq_finish_p            eatq_deal_p 
##                   0.91                   0.97                   0.94 
##     eatq_turn_taking_p    eatq_open_present_p       eatq_before_hw_p 
##                   0.94                   0.91                   0.96 
##     eatq_concentrate_p      eatq_right_away_p      eatq_distracted_p 
##                   0.98                   0.97                   0.95 
##         eatq_impulse_p       eatq_try_focus_p       eatq_finish_hw_p 
##                   0.94                   0.97                   0.96 
##     eatq_early_start_p      eatq_peripheral_p        eatq_puts_off_p 
##                   0.93                   0.96                   0.95 
##   eatq_laugh_control_p     eatq_sidetracked_p   eatq_stick_to_plan_p 
##                   0.92                   0.97                   0.96 
## eatq_close_attention_p 
##                   0.96
```

```
# Bartlett's test for sphericity
cortest.bartlett(data_for_EFA) # highly significant
```

```
## R was not square, finding R from data
```

```
## $chisq
## [1] 63437.46
## 
## $p.value
## [1] 0
## 
## $df
## [1] 378
```

```
# Scree plot to determine the number of factors to extract
scree(data_for_EFA, pc = FALSE)
```

```
# Extract one factor and assess factor loadings
oneF_fit <- factanal(data_for_EFA, 1, rotation = "promax")
oneF_loads <- oneF_fit$loadings
oneF_loads
```

```
## 
## Loadings:
##                        Factor1
## cbcl_q01_p             -0.420 
## cbcl_q04_p             -0.709 
## cbcl_q08_p             -0.768 
## cbcl_q10_p             -0.550 
## cbcl_q13_p             -0.321 
## cbcl_q17_p             -0.436 
## cbcl_q41_p             -0.542 
## cbcl_q61_p             -0.547 
## cbcl_q78_p             -0.755 
## cbcl_q80_p             -0.358 
## eatq_finish_p           0.777 
## eatq_deal_p             0.300 
## eatq_turn_taking_p      0.418 
## eatq_open_present_p     0.291 
## eatq_before_hw_p        0.507 
## eatq_concentrate_p      0.487 
## eatq_right_away_p       0.529 
## eatq_distracted_p       0.509 
## eatq_impulse_p          0.413 
## eatq_try_focus_p        0.639 
## eatq_finish_hw_p        0.652 
## eatq_early_start_p      0.697 
## eatq_peripheral_p       0.666 
## eatq_puts_off_p         0.671 
## eatq_laugh_control_p    0.197 
## eatq_sidetracked_p      0.659 
## eatq_stick_to_plan_p    0.587 
## eatq_close_attention_p  0.617 
## 
##                Factor1
## SS loadings      8.721
## Proportion Var   0.311
```

```
# Create 'ADHD Traits' Bartlett Factor Scores
ADHD_Traits_scores_create <- factanal(data_for_EFA, 1, scores = "Bartlett",
                               rotation = "promax")
ADHD_Traits_scores <- ADHD_Traits_scores_create$scores

# Add factor scores back into table 
comp_data_EFA[, "ADHD_Traits"] <- ADHD_Traits_scores
```

### Create smaller tables with relevant variables

```
# Note - excluding PRS for SCZ and BIP as they showed no association to the ADHD Traits factor scores in PRSice

# ADHD PRS table with ID & pT = .132
data_PRS_ADHD_small <- dplyr::select(data_PRS_ADHD, c('src_subject_id', 'pT_0.132'))
names(data_PRS_ADHD_small)[2] <- 'ADHD_PRS' 

# MDD PRS table with ID & pT = .0945
data_PRS_MDD_small <- dplyr::select(data_PRS_MDD, c('src_subject_id', 'pT_0.0945'))
names(data_PRS_MDD_small)[2] <- 'MDD_PRS' 

# AUT PRS table with ID & pT = .003
data_PRS_AUT_small <- dplyr::select(data_PRS_AUT, c('src_subject_id', 'pT_0.003'))
names(data_PRS_AUT_small)[2] <- 'AUT_PRS' 

# merge all PRS tables
data_PRS <- merge(data_PRS_ADHD_small, data_PRS_MDD_small)
data_PRS <- merge(data_PRS, data_PRS_AUT_small)

# add PCs to the PRS table
data_PRS <- merge(data_PRS, data_PCs)

# ADHD Traits table with ID, ADHD traits, sex, age (change to years), then add age2 and sex*age)
data_ADHD_Traits_small <- dplyr::select(comp_data_EFA, c('src_subject_id', 'interview_age', 'sex', 'ADHD_Traits'))
Age <- (data_ADHD_Traits_small$interview_age)/12
data_ADHD_Traits_small[, "Age"] <- Age
Age2 <- (data_ADHD_Traits_small$Age)^2
data_ADHD_Traits_small[, "Age2"] <- Age2
data_ADHD_Traits_small$Sex <- ifelse(data_ADHD_Traits_small$sex=="M", 1, 2)
AgeSex <- (data_ADHD_Traits_small$Age)*(data_ADHD_Traits_small$Sex)
data_ADHD_Traits_small[, "AgeSex"] <- AgeSex
data_ADHD_Traits_small <- data_ADHD_Traits_small[, c('src_subject_id', 'ADHD_Traits', 'Age', 'Age2', 'Sex', 'AgeSex')]

# WM table with ID & WM
data_WM_2y_small <- dplyr::select(data_WM_2y_QC, c('src_subject_id', 'tfmri_nb_all_beh_c2b_rate'))
# Assess distribution 
hist(data_WM_2y_small$tfmri_nb_all_beh_c2b_rate)
```

```
skewness(data_WM_2y_small$tfmri_nb_all_beh_c2b_rate)
```

```
## [1] -0.6963714
```

```
kurtosis(data_WM_2y_small$tfmri_nb_all_beh_c2b_rate)
```

```
## [1] -0.1255527
```

```
# Apply an arcsine transformation
arc_WM <- asin(sqrt(data_WM_2y_small$tfmri_nb_all_beh_c2b_rate))
data_WM_2y_small[, "arc_WM"] <- arc_WM
hist(data_WM_2y_small$arc_WM)
#Assess the distribution
hist(data_WM_2y_small$arc_WM)
```

```
skewness(data_WM_2y_small$arc_WM)
```

```
## [1] -0.05909523
```

```
kurtosis(data_WM_2y_small$arc_WM)
```

```
## [1] -0.3189293
```

```
# SSRT table with ID & SSRT (and then add a log(SSRT))
data_SSRT_2y_small <- dplyr::select(data_SSRT_2y_QC, c('src_subject_id', 'tfmri_sst_all_beh_total_issrt'))
# Assess distribution 
hist(data_SSRT_2y_small$tfmri_sst_all_beh_total_issrt)
```

```
skewness(data_SSRT_2y_small$tfmri_sst_all_beh_total_issrt)
```

```
## [1] 0.7534291
```

```
kurtosis(data_SSRT_2y_small$tfmri_sst_all_beh_total_issrt)
```

```
## [1] 1.34823
```

```
#Apply an natural log transformation
log_SSRT <- log(data_SSRT_2y_small$tfmri_sst_all_beh_total_issrt)
data_SSRT_2y_small[, "log_SSRT"] <- log_SSRT
#Assess distribution
hist(data_SSRT_2y_small$log_SSRT)
```

```
skewness(data_SSRT_2y_small$log_SSRT)
```

```
## [1] -0.07521215
```

```
kurtosis(data_SSRT_2y_small$log_SSRT)
```

```
## [1] 0.08714305
```

```
# RTV table with ID & RTV (and then add a log(RTV))
data_RTV_2y_small <- dplyr::select(data_RTV_2y_QC, c('src_subject_id', 'tfmri_sst_all_beh_crgo_stdrt'))
# Assess distribution 
hist(data_RTV_2y_small$tfmri_sst_all_beh_crgo_stdrt)
```

```
skewness(data_RTV_2y_small$tfmri_sst_all_beh_crgo_stdrt)
```

```
## [1] -0.09476349
```

```
kurtosis(data_RTV_2y_small$tfmri_sst_all_beh_crgo_stdrt)
```

```
## [1] -0.1794368
```

```
names(data_RTV_2y_small)[2] <- 'RTV'
```

### Create final tables

```
# Join PRS and CBCL
data_final_genbehav <- merge(data_PRS, data_ADHD_Traits_small)

# Final WM table
data_finalWM_group <- merge(data_final_genbehav, data_WM_2y_small)

# Final SSRT table
data_finalSSRT_group <- merge(data_final_genbehav, data_SSRT_2y_small)

# Final RTV table
data_finalRTV_group <- merge(data_final_genbehav, data_RTV_2y_small)
```

### Participant demographics

```
# participants at 2-year follow-up with attention problems according to CBCL
CBCL_clinADHD <- filter(data_CBCL_2y, cbcl_scr_syn_attention_t >= 65)
  # 308/5823 = 5%

# WM group 
mean(data_finalWM_group$Age)
```

```
## [1] 11.93426
```

```
sd(data_finalWM_group$Age)
```

```
## [1] 0.642018
```

```
table(data_finalWM_group$Sex)
```

```
## 
##    1    2 
## 1232  989
```

```
# SSRT group 
mean(data_finalSSRT_group$Age)
```

```
## [1] 11.92548
```

```
sd(data_finalSSRT_group$Age)
```

```
## [1] 0.6422761
```

```
table(data_finalSSRT_group$Sex)
```

```
## 
##    1    2 
## 1095  909
```

```
# RTV group 
mean(data_finalRTV_group$Age)
```

```
## [1] 11.93312
```

```
sd(data_finalRTV_group$Age)
```

```
## [1] 0.6425747
```

```
table(data_finalRTV_group$Sex)
```

```
## 
##    1    2 
## 1165  957
```

### Regressions - cognition regressed onto behaviour

```
# WM - ADHD
regression_WM_ADHD <- lm(ADHD_Traits ~ arc_WM + 
                          Sex + Age + Age2 + AgeSex, data = data_finalWM_group)
summary(regression_WM_ADHD)
```

```
## 
## Call:
## lm(formula = ADHD_Traits ~ arc_WM + Sex + Age + Age2 + AgeSex, 
##     data = data_finalWM_group)
## 
## Residuals:
##     Min      1Q  Median      3Q     Max 
## -3.6838 -0.6051  0.1535  0.7385  2.0102 
## 
## Coefficients:
##              Estimate Std. Error t value Pr(>|t|)    
## (Intercept)  1.484191   8.127849   0.183   0.8551    
## arc_WM       1.545891   0.182609   8.466   <2e-16 ***
## Sex         -1.070207   0.801952  -1.335   0.1822    
## Age         -0.418563   1.349199  -0.310   0.7564    
## Age2         0.007199   0.056187   0.128   0.8981    
## AgeSex       0.124949   0.067149   1.861   0.0629 .  
## ---
## Signif. codes:  0 '***' 0.001 '**' 0.01 '*' 0.05 '.' 0.1 ' ' 1
## 
## Residual standard error: 1.005 on 2215 degrees of freedom
## Multiple R-squared:  0.06736,    Adjusted R-squared:  0.06525 
## F-statistic: 31.99 on 5 and 2215 DF,  p-value: < 2.2e-16
```

```
effectsize::standardize_parameters(regression_WM_ADHD)
```

```
## Registered S3 methods overwritten by 'parameters':
##   method                           from      
##   as.double.parameters_kurtosis    datawizard
##   as.double.parameters_skewness    datawizard
##   as.double.parameters_smoothness  datawizard
##   as.numeric.parameters_kurtosis   datawizard
##   as.numeric.parameters_skewness   datawizard
##   as.numeric.parameters_smoothness datawizard
##   print.parameters_distribution    datawizard
##   print.parameters_kurtosis        datawizard
##   print.parameters_skewness        datawizard
##   summary.parameters_kurtosis      datawizard
##   summary.parameters_skewness      datawizard
```

```
# graph 
visreg(regression_WM_ADHD, xvar = "arc_WM", rug = FALSE, jitter = TRUE, overlay = TRUE, gg = TRUE, line=list(col="#666666"), points=list(col="#66CC66")) + theme_pubr() + labs(x = "arc(Working Memory Function)", y = "ADHD-Like Traits")
```

```
# SSRT - ADHD
regression_SSRT_ADHD <- lm(ADHD_Traits ~ log_SSRT + 
                            Sex + Age + Age2 + AgeSex, data = data_finalSSRT_group)
summary(regression_SSRT_ADHD)
```

```
## 
## Call:
## lm(formula = ADHD_Traits ~ log_SSRT + Sex + Age + Age2 + AgeSex, 
##     data = data_finalSSRT_group)
## 
## Residuals:
##     Min      1Q  Median      3Q     Max 
## -3.5427 -0.6001  0.1504  0.7377  2.0627 
## 
## Coefficients:
##             Estimate Std. Error t value Pr(>|t|)    
## (Intercept)  7.51452    8.51984   0.882    0.378    
## log_SSRT    -0.53091    0.09242  -5.745 1.06e-08 ***
## Sex         -1.12975    0.83840  -1.348    0.178    
## Age         -0.63837    1.41070  -0.453    0.651    
## Age2         0.01752    0.05865   0.299    0.765    
## AgeSex       0.12739    0.07026   1.813    0.070 .  
## ---
## Signif. codes:  0 '***' 0.001 '**' 0.01 '*' 0.05 '.' 0.1 ' ' 1
## 
## Residual standard error: 0.9992 on 1998 degrees of freedom
## Multiple R-squared:  0.05485,    Adjusted R-squared:  0.05249 
## F-statistic: 23.19 on 5 and 1998 DF,  p-value: < 2.2e-16
```

```
effectsize::standardize_parameters(regression_SSRT_ADHD)
```

```
# graph 
visreg(regression_SSRT_ADHD, xvar = "log_SSRT", rug = FALSE, jitter = TRUE, overlay = TRUE, gg = TRUE, line=list(col="#666666"), points=list(col="#339999")) + theme_pubr() + labs(x = "log(Stop Signal Reaction Time)", y = "ADHD-Like Traits")
```

```
# RTV - ADHD
regression_RTV_ADHD <- lm(ADHD_Traits ~ RTV + 
                           Sex + Age + Age2 + AgeSex, data = data_finalRTV_group)
summary(regression_RTV_ADHD)
```

```
## 
## Call:
## lm(formula = ADHD_Traits ~ RTV + Sex + Age + Age2 + AgeSex, data = data_finalRTV_group)
## 
## Residuals:
##     Min      1Q  Median      3Q     Max 
## -3.6283 -0.6137  0.1836  0.7307  2.0571 
## 
## Coefficients:
##               Estimate Std. Error t value Pr(>|t|)    
## (Intercept)  2.8715056  8.3030316   0.346   0.7295    
## RTV         -0.0065023  0.0008544  -7.611 4.08e-14 ***
## Sex         -1.2330599  0.8156540  -1.512   0.1307    
## Age         -0.1843683  1.3775785  -0.134   0.8935    
## Age2        -0.0026904  0.0572837  -0.047   0.9625    
## AgeSex       0.1382039  0.0683134   2.023   0.0432 *  
## ---
## Signif. codes:  0 '***' 0.001 '**' 0.01 '*' 0.05 '.' 0.1 ' ' 1
## 
## Residual standard error: 0.9996 on 2116 degrees of freedom
## Multiple R-squared:  0.0658, Adjusted R-squared:  0.0636 
## F-statistic: 29.81 on 5 and 2116 DF,  p-value: < 2.2e-16
```

```
effectsize::standardize_parameters(regression_RTV_ADHD)
```

```
# graph 
visreg(regression_RTV_ADHD, xvar = "RTV", rug = FALSE, jitter = TRUE, overlay = TRUE, gg = TRUE, line=list(col="#666666"), points=list(col="#6699FF")) + theme_pubr() + labs(x = "Reaction Time Variability", y = "ADHD-Like Traits")
```

```
# Correction for multiple testing 
# Extract p-values from each table
# WM_ADHD
s_regression_WM_ADHD <- summary(regression_WM_ADHD)
p_WM_ADHD <- s_regression_WM_ADHD$coefficients["arc_WM", "Pr(>|t|)"]
# SSRT_ADHD
s_regression_SSRT_ADHD <- summary(regression_SSRT_ADHD)
p_SSRT_ADHD <- s_regression_SSRT_ADHD$coefficients["log_SSRT", "Pr(>|t|)"]
# RTV_ADHD
s_regression_RTV_ADHD <- summary(regression_RTV_ADHD)
p_RTV_ADHD <- s_regression_RTV_ADHD$coefficients["RTV", "Pr(>|t|)"]
# Create a vector of p-values
corr_cog_beh <- c(p_WM_ADHD, p_SSRT_ADHD, p_RTV_ADHD)
# Apply bonferroni correction
p.adjust(corr_cog_beh, method = "bonferroni") # all significant
```

```
## [1] 1.370402e-16 3.189053e-08 1.224591e-13
```

### Regressions - ADHD PRS regressed onto cognition

```
# ADHD PRS - WM
regression_ADHD_PRS_WM <- lm(formula = arc_WM ~ ADHD_PRS + 
                          Sex + Age + Age2 + AgeSex + PC1 + PC2 + PC3, data = data_finalWM_group)
summary(regression_ADHD_PRS_WM)
```

```
## 
## Call:
## lm(formula = arc_WM ~ ADHD_PRS + Sex + Age + Age2 + AgeSex + 
##     PC1 + PC2 + PC3, data = data_finalWM_group)
## 
## Residuals:
##      Min       1Q   Median       3Q      Max 
## -0.39029 -0.07435  0.00737  0.07943  0.35379 
## 
## Coefficients:
##               Estimate Std. Error t value Pr(>|t|)    
## (Intercept)  1.893e+00  9.413e-01   2.011   0.0445 *  
## ADHD_PRS    -2.620e+02  5.288e+01  -4.955 7.78e-07 ***
## Sex         -4.651e-02  9.288e-02  -0.501   0.6166    
## Age         -1.317e-01  1.563e-01  -0.843   0.3994    
## Age2         6.972e-03  6.509e-03   1.071   0.2843    
## AgeSex       2.573e-03  7.778e-03   0.331   0.7408    
## PC1         -9.702e-01  1.394e+00  -0.696   0.4867    
## PC2          6.810e-02  1.678e+00   0.041   0.9676    
## PC3         -1.160e+00  1.091e+00  -1.063   0.2878    
## ---
## Signif. codes:  0 '***' 0.001 '**' 0.01 '*' 0.05 '.' 0.1 ' ' 1
## 
## Residual standard error: 0.1163 on 2212 degrees of freedom
## Multiple R-squared:  0.05983,    Adjusted R-squared:  0.05643 
## F-statistic:  17.6 on 8 and 2212 DF,  p-value: < 2.2e-16
```

```
effectsize::standardize_parameters(regression_ADHD_PRS_WM)
```

```
# graph 
visreg(regression_ADHD_PRS_WM, xvar = "ADHD_PRS", rug = FALSE, jitter = TRUE, overlay = TRUE, gg = TRUE, line=list(col="#666666"), points=list(col="#66CC66")) + theme_pubr() + labs(x = "ADHD PRS", y = "arc(Working Memory Function)")
```

```
# ADHD PRS - SSRT
regression_ADHD_PRS_SSRT <- lm(formula = log_SSRT ~ ADHD_PRS +
                            Sex + Age + Age2 + AgeSex + PC1 + PC2 + PC3, data = data_finalSSRT_group)
summary(regression_ADHD_PRS_SSRT)
```

```
## 
## Call:
## lm(formula = log_SSRT ~ ADHD_PRS + Sex + Age + Age2 + AgeSex + 
##     PC1 + PC2 + PC3, data = data_finalSSRT_group)
## 
## Residuals:
##      Min       1Q   Median       3Q      Max 
## -0.72037 -0.15908  0.00456  0.16191  0.91007 
## 
## Coefficients:
##               Estimate Std. Error t value Pr(>|t|)   
## (Intercept)   5.654676   2.061173   2.743  0.00613 **
## ADHD_PRS    234.929573 116.522361   2.016  0.04392 * 
## Sex          -0.123574   0.202966  -0.609  0.54270   
## Age           0.019286   0.341780   0.056  0.95501   
## Age2         -0.002696   0.014211  -0.190  0.84954   
## AgeSex        0.008674   0.017011   0.510  0.61016   
## PC1           2.276855   3.037419   0.750  0.45358   
## PC2          -2.687410   3.705798  -0.725  0.46842   
## PC3          -0.005208   2.382833  -0.002  0.99826   
## ---
## Signif. codes:  0 '***' 0.001 '**' 0.01 '*' 0.05 '.' 0.1 ' ' 1
## 
## Residual standard error: 0.2417 on 1995 degrees of freedom
## Multiple R-squared:  0.01139,    Adjusted R-squared:  0.00743 
## F-statistic: 2.874 on 8 and 1995 DF,  p-value: 0.0035
```

```
effectsize::standardize_parameters(regression_ADHD_PRS_SSRT)
```

```
# graph 
visreg(regression_ADHD_PRS_SSRT, xvar = "ADHD_PRS", rug = FALSE, jitter = TRUE, overlay = TRUE, gg = TRUE, line=list(col="#666666"), points=list(col="#339999")) +
  theme_pubr() +
  labs(x = "ADHD PRS", y = "log(Stop Signal Reaction Time)")
```

```
# ADHD PRS - RTV
regression_ADHD_PRS_RTV <- lm(formula = RTV ~ ADHD_PRS + 
                           Sex + Age + Age2 + AgeSex + PC1 + PC2 + PC3, data = data_finalRTV_group)
summary(regression_ADHD_PRS_RTV)
```

```
## 
## Call:
## lm(formula = RTV ~ ADHD_PRS + Sex + Age + Age2 + AgeSex + PC1 + 
##     PC2 + PC3, data = data_finalRTV_group)
## 
## Residuals:
##     Min      1Q  Median      3Q     Max 
## -70.187 -17.204   0.667  16.563  94.053 
## 
## Coefficients:
##              Estimate Std. Error t value Pr(>|t|)    
## (Intercept)    36.423    209.463   0.174 0.861971    
## ADHD_PRS    67838.223  11847.216   5.726 1.17e-08 ***
## Sex            -5.023     20.549  -0.244 0.806914    
## Age            20.047     34.739   0.577 0.563944    
## Age2           -1.151      1.444  -0.797 0.425710    
## AgeSex          0.488      1.721   0.284 0.776779    
## PC1            49.077    308.655   0.159 0.873681    
## PC2          -480.926    374.343  -1.285 0.199031    
## PC3           841.029    241.480   3.483 0.000506 ***
## ---
## Signif. codes:  0 '***' 0.001 '**' 0.01 '*' 0.05 '.' 0.1 ' ' 1
## 
## Residual standard error: 25.17 on 2113 degrees of freedom
## Multiple R-squared:  0.05005,    Adjusted R-squared:  0.04645 
## F-statistic: 13.92 on 8 and 2113 DF,  p-value: < 2.2e-16
```

```
effectsize::standardize_parameters(regression_ADHD_PRS_RTV)
```

```
# graph 
visreg(regression_ADHD_PRS_RTV, xvar = "ADHD_PRS", rug = FALSE, jitter = TRUE, overlay = TRUE, gg = TRUE, line=list(col="#666666"), points=list(col="#6699FF")) +
  theme_pubr() +
  labs(x = "ADHD PRS", y = "Reaction Time Variability")
```

```
# Correction for multiple testing 
# Extract p-values from each table
# ADHD_PRS_WM
s_regression_ADHD_PRS_WM <- summary(regression_ADHD_PRS_WM)
p_ADHD_PRS_WM <- s_regression_ADHD_PRS_WM$coefficients["ADHD_PRS", "Pr(>|t|)"]
# ADHD_PRS_SSRT
s_regression_ADHD_PRS_SSRT <- summary(regression_ADHD_PRS_SSRT)
p_ADHD_PRS_SSRT <- s_regression_ADHD_PRS_SSRT$coefficients["ADHD_PRS", "Pr(>|t|)"]
# ADHD_PRS_RTV
s_regression_ADHD_PRS_RTV <- summary(regression_ADHD_PRS_RTV)
p_ADHD_PRS_RTV <- s_regression_ADHD_PRS_RTV$coefficients["ADHD_PRS", "Pr(>|t|)"]
# Create a vector of p-values
corr_gen_cog <- c(p_ADHD_PRS_WM, p_ADHD_PRS_SSRT, p_ADHD_PRS_RTV)
# Apply bonferroni correction
p.adjust(corr_gen_cog, method = "bonferroni") # WM and RTV significant
```

```
## [1] 2.333895e-06 1.317462e-01 3.523493e-08
```

### Regressions - AUT PRS & MDD PRS regressed onto cognition

```
# AUT PRS - WM
regression_AUT_PRS_WM <- lm(formula = arc_WM ~ AUT_PRS + 
                          Sex + Age + Age2 + AgeSex + PC1 + PC2 + PC3, data = data_finalWM_group)
summary(regression_AUT_PRS_WM)
```

```
## 
## Call:
## lm(formula = arc_WM ~ AUT_PRS + Sex + Age + Age2 + AgeSex + PC1 + 
##     PC2 + PC3, data = data_finalWM_group)
## 
## Residuals:
##      Min       1Q   Median       3Q      Max 
## -0.36739 -0.07429  0.00743  0.07916  0.35680 
## 
## Coefficients:
##              Estimate Std. Error t value Pr(>|t|)  
## (Intercept)  1.913059   0.947218   2.020   0.0435 *
## AUT_PRS     -5.059243   7.226279  -0.700   0.4839  
## Sex         -0.039459   0.093381  -0.423   0.6727  
## Age         -0.155669   0.157145  -0.991   0.3220  
## Age2         0.008003   0.006544   1.223   0.2215  
## AgeSex       0.001905   0.007819   0.244   0.8075  
## PC1         -1.319244   1.401063  -0.942   0.3465  
## PC2         -0.360364   1.685299  -0.214   0.8307  
## PC3         -0.893772   1.096983  -0.815   0.4153  
## ---
## Signif. codes:  0 '***' 0.001 '**' 0.01 '*' 0.05 '.' 0.1 ' ' 1
## 
## Residual standard error: 0.1169 on 2212 degrees of freedom
## Multiple R-squared:  0.04961,    Adjusted R-squared:  0.04617 
## F-statistic: 14.43 on 8 and 2212 DF,  p-value: < 2.2e-16
```

```
effectsize::standardize_parameters(regression_AUT_PRS_WM)
```

```
# graph 
visreg(regression_AUT_PRS_WM, xvar = "AUT_PRS", rug = FALSE, jitter = TRUE, overlay = TRUE, gg = TRUE, line=list(col="#666666"), points=list(col="#66CC66")) +
  theme_pubr() +
  labs(x = "AUT PRS", y = "Working Memory Function")
```

```
# MDD PRS - WM
regression_MDD_PRS_WM <- lm(formula = arc_WM ~ MDD_PRS + 
                          Sex + Age + Age2 + AgeSex + PC1 + PC2 + PC3, data = data_finalWM_group)
summary(regression_MDD_PRS_WM)
```

```
## 
## Call:
## lm(formula = arc_WM ~ MDD_PRS + Sex + Age + Age2 + AgeSex + PC1 + 
##     PC2 + PC3, data = data_finalWM_group)
## 
## Residuals:
##      Min       1Q   Median       3Q      Max 
## -0.36915 -0.07423  0.00803  0.07906  0.36048 
## 
## Coefficients:
##               Estimate Std. Error t value Pr(>|t|)  
## (Intercept)  1.845e+00  9.484e-01   1.946   0.0518 .
## MDD_PRS     -1.514e+02  1.058e+02  -1.432   0.1524  
## Sex         -4.672e-02  9.345e-02  -0.500   0.6171  
## Age         -1.494e-01  1.571e-01  -0.951   0.3416  
## Age2         7.705e-03  6.541e-03   1.178   0.2389  
## AgeSex       2.477e-03  7.824e-03   0.317   0.7516  
## PC1         -1.234e+00  1.401e+00  -0.881   0.3785  
## PC2         -3.680e-01  1.685e+00  -0.218   0.8271  
## PC3         -8.351e-01  1.095e+00  -0.763   0.4458  
## ---
## Signif. codes:  0 '***' 0.001 '**' 0.01 '*' 0.05 '.' 0.1 ' ' 1
## 
## Residual standard error: 0.1169 on 2212 degrees of freedom
## Multiple R-squared:  0.05028,    Adjusted R-squared:  0.04684 
## F-statistic: 14.64 on 8 and 2212 DF,  p-value: < 2.2e-16
```

```
effectsize::standardize_parameters(regression_MDD_PRS_WM)
```

```
# graph 
visreg(regression_MDD_PRS_WM, xvar = "MDD_PRS", rug = FALSE, jitter = TRUE, overlay = TRUE, gg = TRUE, line=list(col="#666666"), points=list(col="#66CC66")) +
  theme_pubr() +
  labs(x = "MDD PRS", y = "Working Memory Function")
```

```
# AUT PRS - SSRT
regression_AUT_PRS_SSRT <- lm(formula = log_SSRT ~ AUT_PRS +
                            Sex + Age + Age2 + AgeSex + PC1 + PC2 + PC3, data = data_finalSSRT_group)
summary(regression_AUT_PRS_SSRT)
```

```
## 
## Call:
## lm(formula = log_SSRT ~ AUT_PRS + Sex + Age + Age2 + AgeSex + 
##     PC1 + PC2 + PC3, data = data_finalSSRT_group)
## 
## Residuals:
##     Min      1Q  Median      3Q     Max 
## -0.7270 -0.1587  0.0035  0.1626  0.9180 
## 
## Coefficients:
##               Estimate Std. Error t value Pr(>|t|)   
## (Intercept)   5.481895   2.063995   2.656  0.00797 **
## AUT_PRS     -20.080397  15.627508  -1.285  0.19896   
## Sex          -0.124676   0.203101  -0.614  0.53938   
## Age           0.032086   0.341887   0.094  0.92524   
## Age2         -0.003237   0.014215  -0.228  0.81989   
## AgeSex        0.008940   0.017020   0.525  0.59948   
## PC1           2.503211   3.035808   0.825  0.40972   
## PC2          -2.425443   3.704511  -0.655  0.51272   
## PC3          -0.469279   2.386267  -0.197  0.84412   
## ---
## Signif. codes:  0 '***' 0.001 '**' 0.01 '*' 0.05 '.' 0.1 ' ' 1
## 
## Residual standard error: 0.2419 on 1995 degrees of freedom
## Multiple R-squared:  0.0102, Adjusted R-squared:  0.00623 
## F-statistic:  2.57 on 8 and 1995 DF,  p-value: 0.008657
```

```
effectsize::standardize_parameters(regression_AUT_PRS_SSRT)
```

```
# graph 
visreg(regression_AUT_PRS_SSRT, xvar = "AUT_PRS", rug = FALSE, jitter = TRUE, overlay = TRUE, gg = TRUE, line=list(col="#666666"), points=list(col="#339999")) +
  theme_pubr() +
  labs(x = "AUT PRS", y = "Response Inhibition")
```

```
# MDD PRS - SSRT
regression_MDD_PRS_SSRT <- lm(formula = log_SSRT ~ MDD_PRS +
                            Sex + Age + Age2 + AgeSex + PC1 + PC2 + PC3, data = data_finalSSRT_group)
summary(regression_MDD_PRS_SSRT)
```

```
## 
## Call:
## lm(formula = log_SSRT ~ MDD_PRS + Sex + Age + Age2 + AgeSex + 
##     PC1 + PC2 + PC3, data = data_finalSSRT_group)
## 
## Residuals:
##      Min       1Q   Median       3Q      Max 
## -0.72595 -0.15729  0.00578  0.16289  0.92791 
## 
## Coefficients:
##               Estimate Std. Error t value Pr(>|t|)   
## (Intercept)   5.771845   2.068916   2.790  0.00532 **
## MDD_PRS     250.008592 228.647957   1.093  0.27434   
## Sex          -0.119059   0.203308  -0.586  0.55821   
## Age           0.031969   0.341961   0.093  0.92553   
## Age2         -0.003202   0.014219  -0.225  0.82185   
## AgeSex        0.008448   0.017037   0.496  0.62003   
## PC1           2.551037   3.035417   0.840  0.40077   
## PC2          -2.309102   3.704174  -0.623  0.53311   
## PC3          -0.272652   2.381135  -0.115  0.90885   
## ---
## Signif. codes:  0 '***' 0.001 '**' 0.01 '*' 0.05 '.' 0.1 ' ' 1
## 
## Residual standard error: 0.2419 on 1995 degrees of freedom
## Multiple R-squared:  0.009973,   Adjusted R-squared:  0.006003 
## F-statistic: 2.512 on 8 and 1995 DF,  p-value: 0.01024
```

```
effectsize::standardize_parameters(regression_MDD_PRS_SSRT)
```

```
# graph 
visreg(regression_MDD_PRS_SSRT, xvar = "MDD_PRS", rug = FALSE, jitter = TRUE, overlay = TRUE, gg = TRUE, line=list(col="#666666"), points=list(col="#339999")) +
  theme_pubr() +
  labs(x = "MMD PRS", y = "Response Inhibition")
```

```
# AUT PRS - RTV
regression_AUT_PRS_RTV <- lm(formula = RTV ~ AUT_PRS + 
                           Sex + Age + Age2 + AgeSex + PC1 + PC2 + PC3, data = data_finalRTV_group)
summary(regression_AUT_PRS_RTV)
```

```
## 
## Call:
## lm(formula = RTV ~ AUT_PRS + Sex + Age + Age2 + AgeSex + PC1 + 
##     PC2 + PC3, data = data_finalRTV_group)
## 
## Residuals:
##     Min      1Q  Median      3Q     Max 
## -70.356 -17.430   0.761  17.322  95.485 
## 
## Coefficients:
##              Estimate Std. Error t value Pr(>|t|)   
## (Intercept)   18.2498   211.1790   0.086  0.93114   
## AUT_PRS      647.0931  1605.0385   0.403  0.68687   
## Sex           -5.9429    20.7083  -0.287  0.77415   
## Age           27.4213    34.9982   0.784  0.43342   
## Age2          -1.4643     1.4553  -1.006  0.31443   
## AgeSex         0.5936     1.7343   0.342  0.73220   
## PC1          147.5590   310.6978   0.475  0.63489   
## PC2         -370.9201   376.8337  -0.984  0.32508   
## PC3          776.1168   243.5353   3.187  0.00146 **
## ---
## Signif. codes:  0 '***' 0.001 '**' 0.01 '*' 0.05 '.' 0.1 ' ' 1
## 
## Residual standard error: 25.37 on 2113 degrees of freedom
## Multiple R-squared:  0.03538,    Adjusted R-squared:  0.03173 
## F-statistic: 9.689 on 8 and 2113 DF,  p-value: 2.798e-13
```

```
effectsize::standardize_parameters(regression_AUT_PRS_RTV)
```

```
# graph 
visreg(regression_AUT_PRS_RTV, xvar = "AUT_PRS", rug = FALSE, jitter = TRUE, overlay = TRUE, gg = TRUE, line=list(col="#666666"), points=list(col="#6699FF")) +
  theme_pubr() +
  labs(x = "AUT PRS", y = "Reaction Time Variability")
```

```
# MDD PRS - RTV
regression_MDD_PRS_RTV <- lm(formula = RTV ~ MDD_PRS + 
                           Sex + Age + Age2 + AgeSex + PC1 + PC2 + PC3, data = data_finalRTV_group)
summary(regression_MDD_PRS_RTV)
```

```
## 
## Call:
## lm(formula = RTV ~ MDD_PRS + Sex + Age + Age2 + AgeSex + PC1 + 
##     PC2 + PC3, data = data_finalRTV_group)
## 
## Residuals:
##     Min      1Q  Median      3Q     Max 
## -69.973 -17.191   0.841  17.092  95.119 
## 
## Coefficients:
##               Estimate Std. Error t value Pr(>|t|)   
## (Intercept)    55.0660   211.5139   0.260  0.79463   
## MDD_PRS     53051.3779 23261.3437   2.281  0.02267 * 
## Sex            -3.5755    20.7051  -0.173  0.86291   
## Age            24.6214    34.9592   0.704  0.48133   
## Age2           -1.3358     1.4538  -0.919  0.35828   
## AgeSex          0.4046     1.7340   0.233  0.81553   
## PC1           123.5961   310.3393   0.398  0.69048   
## PC2          -366.0114   376.3212  -0.973  0.33086   
## PC3           765.9162   242.7289   3.155  0.00163 **
## ---
## Signif. codes:  0 '***' 0.001 '**' 0.01 '*' 0.05 '.' 0.1 ' ' 1
## 
## Residual standard error: 25.34 on 2113 degrees of freedom
## Multiple R-squared:  0.03768,    Adjusted R-squared:  0.03404 
## F-statistic: 10.34 on 8 and 2113 DF,  p-value: 2.715e-14
```

```
effectsize::standardize_parameters(regression_MDD_PRS_RTV)
```

```
# graph 
visreg(regression_MDD_PRS_RTV, xvar = "MDD_PRS", rug = FALSE, jitter = TRUE, overlay = TRUE, gg = TRUE, line=list(col="#666666"), points=list(col="#6699FF")) +
  theme_pubr() +
  labs(x = "MDD PRS", y = "Reaction Time Variability")
```

```
# Correction for multiple testing 
# Extract p-values from each table
# AUT_PRS_WM
s_regression_AUT_PRS_WM <- summary(regression_AUT_PRS_WM)
p_AUT_PRS_WM <- s_regression_AUT_PRS_WM$coefficients["AUT_PRS", "Pr(>|t|)"]
# MDD_PRS_WM
s_regression_MDD_PRS_WM <- summary(regression_MDD_PRS_WM)
p_MDD_PRS_WM <- s_regression_MDD_PRS_WM$coefficients["MDD_PRS", "Pr(>|t|)"]
# AUT_PRS_SSRT
s_regression_AUT_PRS_SSRT <- summary(regression_AUT_PRS_SSRT)
p_AUT_PRS_SSRT <- s_regression_AUT_PRS_SSRT$coefficients["AUT_PRS", "Pr(>|t|)"]
# MDD_PRS_SSRT
s_regression_MDD_PRS_SSRT <- summary(regression_MDD_PRS_SSRT)
p_MDD_PRS_SSRT <- s_regression_MDD_PRS_SSRT$coefficients["MDD_PRS", "Pr(>|t|)"]
# AUT_PRS_RTV
s_regression_AUT_PRS_RTV <- summary(regression_AUT_PRS_RTV)
p_AUT_PRS_RTV <- s_regression_AUT_PRS_RTV$coefficients["AUT_PRS", "Pr(>|t|)"]
# MDD_PRS_RTV
s_regression_MDD_PRS_RTV <- summary(regression_MDD_PRS_RTV)
p_MDD_PRS_RTV <- s_regression_MDD_PRS_RTV$coefficients["MDD_PRS", "Pr(>|t|)"]
# Create a vector of p-values
corr_spec <- c(p_AUT_PRS_WM, p_MDD_PRS_WM, p_AUT_PRS_SSRT, p_MDD_PRS_SSRT,
                 p_AUT_PRS_RTV, p_MDD_PRS_RTV)
# Apply bonferroni correction
p.adjust(corr_spec, method = "bonferroni") # none are significant
```

```
## [1] 1.0000000 0.9143127 1.0000000 1.0000000 1.0000000 0.1360039
```

### Mediation - ADHD PRS | working memory | ADHD traits

```
#Set a seed for the sake of replicability when bootstrapping
set.seed(5)

# Create the model
WM_mediation_model <- mediation1(y = "ADHD_Traits", #DV
                     x = "ADHD_PRS", #IV 
                     m = "arc_WM", #Mediator
                     cvs = c("Age", "Sex", "Age2", "AgeSex", "PC1", "PC2", "PC3"), #Covariates
                     df = data_finalWM_group, #Datafram
                     with_out = T, #Including outliers
                     nboot = 5000, #Number of bootstraps
                     conf_level = .95 #CI width
                     )
```

```
#bootstrapped indirect effect 
WM_mediation_model$boot.results
```

```
## 
## ORDINARY NONPARAMETRIC BOOTSTRAP
## 
## 
## Call:
## boot(data = finaldata, statistic = indirectmed, R = nboot, formula2 = allformulas$eq2, 
##     formula3 = allformulas$eq3, x = x, med.var = m)
## 
## 
## Bootstrap Statistics :
##     original    bias    std. error
## t1* -378.701 -1.689526    92.15222
```

```
#bootstrapped CI (test of significance)
WM_mediation_model$boot.ci # doesn't include 0 = significant
```

```
## BOOTSTRAP CONFIDENCE INTERVAL CALCULATIONS
## Based on 5000 bootstrap replicates
## 
## CALL : 
## boot.ci(boot.out = bootresults, conf = conf_level, type = "norm")
## 
## Intervals : 
## Level      Normal        
## 95%   (-557.6, -196.4 )  
## Calculations and Intervals on Original Scale
```

```
# STANDARDIZE results
# fit the models
# a path
mediation_1 <- lm(arc_WM ~ ADHD_PRS + Sex + Age + Age2 + AgeSex + PC1 + PC2 + PC3, 
                  data = data_finalWM_group)
summary(mediation_1)
```

```
## 
## Call:
## lm(formula = arc_WM ~ ADHD_PRS + Sex + Age + Age2 + AgeSex + 
##     PC1 + PC2 + PC3, data = data_finalWM_group)
## 
## Residuals:
##      Min       1Q   Median       3Q      Max 
## -0.39029 -0.07435  0.00737  0.07943  0.35379 
## 
## Coefficients:
##               Estimate Std. Error t value Pr(>|t|)    
## (Intercept)  1.893e+00  9.413e-01   2.011   0.0445 *  
## ADHD_PRS    -2.620e+02  5.288e+01  -4.955 7.78e-07 ***
## Sex         -4.651e-02  9.288e-02  -0.501   0.6166    
## Age         -1.317e-01  1.563e-01  -0.843   0.3994    
## Age2         6.972e-03  6.509e-03   1.071   0.2843    
## AgeSex       2.573e-03  7.778e-03   0.331   0.7408    
## PC1         -9.702e-01  1.394e+00  -0.696   0.4867    
## PC2          6.810e-02  1.678e+00   0.041   0.9676    
## PC3         -1.160e+00  1.091e+00  -1.063   0.2878    
## ---
## Signif. codes:  0 '***' 0.001 '**' 0.01 '*' 0.05 '.' 0.1 ' ' 1
## 
## Residual standard error: 0.1163 on 2212 degrees of freedom
## Multiple R-squared:  0.05983,    Adjusted R-squared:  0.05643 
## F-statistic:  17.6 on 8 and 2212 DF,  p-value: < 2.2e-16
```

```
effectsize::standardize_parameters(mediation_1)
```

```
# b and c' paths
mediation_2 <- lm(ADHD_Traits ~ ADHD_PRS + arc_WM + Sex + Age + Age2 + AgeSex + PC1 + PC2 + PC3, 
                  data = data_finalWM_group)
summary(mediation_2)
```

```
## 
## Call:
## lm(formula = ADHD_Traits ~ ADHD_PRS + arc_WM + Sex + Age + Age2 + 
##     AgeSex + PC1 + PC2 + PC3, data = data_finalWM_group)
## 
## Residuals:
##     Min      1Q  Median      3Q     Max 
## -3.5472 -0.6125  0.1272  0.7384  2.1350 
## 
## Coefficients:
##               Estimate Std. Error t value Pr(>|t|)    
## (Intercept)  1.180e+00  8.096e+00   0.146   0.8841    
## ADHD_PRS    -2.374e+03  4.569e+02  -5.195 2.23e-07 ***
## arc_WM       1.445e+00  1.827e-01   7.910 4.03e-15 ***
## Sex         -1.129e+00  7.982e-01  -1.415   0.1573    
## Age         -2.354e-01  1.343e+00  -0.175   0.8609    
## Age2        -5.029e-04  5.595e-02  -0.009   0.9928    
## AgeSex       1.306e-01  6.683e-02   1.953   0.0509 .  
## PC1         -3.515e-01  1.198e+01  -0.029   0.9766    
## PC2          6.369e+00  1.442e+01   0.442   0.6587    
## PC3         -6.776e+00  9.379e+00  -0.723   0.4701    
## ---
## Signif. codes:  0 '***' 0.001 '**' 0.01 '*' 0.05 '.' 0.1 ' ' 1
## 
## Residual standard error: 0.9995 on 2211 degrees of freedom
## Multiple R-squared:  0.0787, Adjusted R-squared:  0.07495 
## F-statistic: 20.99 on 9 and 2211 DF,  p-value: < 2.2e-16
```

```
effectsize::standardize_parameters(mediation_2)
```

```
# c path
mediation_3 <- lm(ADHD_Traits ~ ADHD_PRS + Sex + Age + Age2 + AgeSex + PC1 + PC2 + PC3, 
                  data = data_finalWM_group)
summary(mediation_3)
```

```
## 
## Call:
## lm(formula = ADHD_Traits ~ ADHD_PRS + Sex + Age + Age2 + AgeSex + 
##     PC1 + PC2 + PC3, data = data_finalWM_group)
## 
## Residuals:
##     Min      1Q  Median      3Q     Max 
## -3.5023 -0.6172  0.1371  0.7520  2.0681 
## 
## Coefficients:
##               Estimate Std. Error t value Pr(>|t|)    
## (Intercept)  3.916e+00  8.201e+00   0.478   0.6330    
## ADHD_PRS    -2.753e+03  4.607e+02  -5.975 2.68e-09 ***
## Sex         -1.196e+00  8.091e-01  -1.479   0.1394    
## Age         -4.258e-01  1.362e+00  -0.313   0.7546    
## Age2         9.573e-03  5.671e-02   0.169   0.8660    
## AgeSex       1.343e-01  6.776e-02   1.982   0.0476 *  
## PC1         -1.754e+00  1.215e+01  -0.144   0.8852    
## PC2          6.467e+00  1.462e+01   0.442   0.6582    
## PC3         -8.453e+00  9.506e+00  -0.889   0.3740    
## ---
## Signif. codes:  0 '***' 0.001 '**' 0.01 '*' 0.05 '.' 0.1 ' ' 1
## 
## Residual standard error: 1.013 on 2212 degrees of freedom
## Multiple R-squared:  0.05263,    Adjusted R-squared:  0.0492 
## F-statistic: 15.36 on 8 and 2212 DF,  p-value: < 2.2e-16
```

```
effectsize::standardize_parameters(mediation_3)
```

```
# standardized indirect effect
WM_M1 <- lm.beta(mediation_1) # standardized a path
WM_M2 <- lm.beta(mediation_2) # standardized b path
WM_ie <- (WM_M1$standardized.coefficients["ADHD_PRS"])*(WM_M2$standardized.coefficients["arc_WM"]) # times the standardized a path by the standardized b path
WM_ie
```

```
##    ADHD_PRS 
## -0.01716507
```

```
# proportion mediated
WM_M3 <- lm.beta(mediation_3) # standardized c (total) path
WM_ie/(WM_M3$standardized.coefficients["ADHD_PRS"])# divide the standardized indirect effect by the standardized total effect
```

```
##  ADHD_PRS 
## 0.1375818
```

### Mediation - ADHD PRS | reaction time variability | ADHD traits

```
#Set a seed for the sake of replicability when bootstrapping
set.seed(5)

# Create the model
RTV_mediation_model <- mediation1(y = "ADHD_Traits", #DV
                     x = "ADHD_PRS", #IV 
                     m = "RTV", #Mediator
                     cvs = c("Age", "Sex", "Age2", "AgeSex", "PC1", "PC2", "PC3"), #Covariates
                     df = data_finalRTV_group, #Datafram
                     with_out = T, #Including outliers
                     nboot = 5000, #Number of bootstraps
                     conf_level = .95 #CI width
                     )
```

```
#bootstrapped indirect effect 
RTV_mediation_model$boot.results
```

```
## 
## ORDINARY NONPARAMETRIC BOOTSTRAP
## 
## 
## Call:
## boot(data = finaldata, statistic = indirectmed, R = nboot, formula2 = allformulas$eq2, 
##     formula3 = allformulas$eq3, x = x, med.var = m)
## 
## 
## Bootstrap Statistics :
##      original   bias    std. error
## t1* -412.1706 2.930065    91.64532
```

```
#bootstrapped CI (test of significance)
RTV_mediation_model$boot.ci # doesn't include 0 = significant
```

```
## BOOTSTRAP CONFIDENCE INTERVAL CALCULATIONS
## Based on 5000 bootstrap replicates
## 
## CALL : 
## boot.ci(boot.out = bootresults, conf = conf_level, type = "norm")
## 
## Intervals : 
## Level      Normal        
## 95%   (-594.7, -235.5 )  
## Calculations and Intervals on Original Scale
```

```
# STANDARDIZE results
# fit the modelss

# a path
mediation_4 <- lm(RTV ~ ADHD_PRS + Sex + Age + Age2 + AgeSex + PC1 + PC2 + PC3, 
                  data = data_finalRTV_group)
summary(mediation_4)
```

```
## 
## Call:
## lm(formula = RTV ~ ADHD_PRS + Sex + Age + Age2 + AgeSex + PC1 + 
##     PC2 + PC3, data = data_finalRTV_group)
## 
## Residuals:
##     Min      1Q  Median      3Q     Max 
## -70.187 -17.204   0.667  16.563  94.053 
## 
## Coefficients:
##              Estimate Std. Error t value Pr(>|t|)    
## (Intercept)    36.423    209.463   0.174 0.861971    
## ADHD_PRS    67838.223  11847.216   5.726 1.17e-08 ***
## Sex            -5.023     20.549  -0.244 0.806914    
## Age            20.047     34.739   0.577 0.563944    
## Age2           -1.151      1.444  -0.797 0.425710    
## AgeSex          0.488      1.721   0.284 0.776779    
## PC1            49.077    308.655   0.159 0.873681    
## PC2          -480.926    374.343  -1.285 0.199031    
## PC3           841.029    241.480   3.483 0.000506 ***
## ---
## Signif. codes:  0 '***' 0.001 '**' 0.01 '*' 0.05 '.' 0.1 ' ' 1
## 
## Residual standard error: 25.17 on 2113 degrees of freedom
## Multiple R-squared:  0.05005,    Adjusted R-squared:  0.04645 
## F-statistic: 13.92 on 8 and 2113 DF,  p-value: < 2.2e-16
```

```
effectsize::standardize_parameters(mediation_4)
```

```
# b and c' paths
mediation_5 <- lm(ADHD_Traits ~ ADHD_PRS + RTV + Sex + Age + Age2 + AgeSex + PC1 + PC2 + PC3, 
                  data = data_finalRTV_group)
summary(mediation_5)
```

```
## 
## Call:
## lm(formula = ADHD_Traits ~ ADHD_PRS + RTV + Sex + Age + Age2 + 
##     AgeSex + PC1 + PC2 + PC3, data = data_finalRTV_group)
## 
## Residuals:
##     Min      1Q  Median      3Q     Max 
## -3.5238 -0.6041  0.1460  0.7296  2.1301 
## 
## Coefficients:
##               Estimate Std. Error t value Pr(>|t|)    
## (Intercept)  2.219e+00  8.279e+00   0.268   0.7887    
## ADHD_PRS    -2.243e+03  4.719e+02  -4.753 2.14e-06 ***
## RTV         -6.076e-03  8.598e-04  -7.066 2.16e-12 ***
## Sex         -1.261e+00  8.122e-01  -1.552   0.1208    
## Age          3.590e-02  1.373e+00   0.026   0.9791    
## Age2        -1.199e-02  5.710e-02  -0.210   0.8337    
## AgeSex       1.414e-01  6.803e-02   2.079   0.0378 *  
## PC1          7.712e+00  1.220e+01   0.632   0.5273    
## PC2         -3.207e+00  1.480e+01  -0.217   0.8285    
## PC3          6.621e+00  9.572e+00   0.692   0.4892    
## ---
## Signif. codes:  0 '***' 0.001 '**' 0.01 '*' 0.05 '.' 0.1 ' ' 1
## 
## Residual standard error: 0.9949 on 2112 degrees of freedom
## Multiple R-squared:  0.07628,    Adjusted R-squared:  0.07234 
## F-statistic: 19.38 on 9 and 2112 DF,  p-value: < 2.2e-16
```

```
effectsize::standardize_parameters(mediation_5)
```

```
# c path
mediation_6 <- lm(ADHD_Traits ~ ADHD_PRS + Sex + Age + Age2 + AgeSex + PC1 + PC2 + PC3, 
                  data = data_finalRTV_group)
summary(mediation_6)
```

```
## 
## Call:
## lm(formula = ADHD_Traits ~ ADHD_PRS + Sex + Age + Age2 + AgeSex + 
##     PC1 + PC2 + PC3, data = data_finalRTV_group)
## 
## Residuals:
##     Min      1Q  Median      3Q     Max 
## -3.5304 -0.6174  0.1403  0.7455  2.0590 
## 
## Coefficients:
##               Estimate Std. Error t value Pr(>|t|)    
## (Intercept)  1.997e+00  8.374e+00   0.239   0.8115    
## ADHD_PRS    -2.655e+03  4.737e+02  -5.605 2.35e-08 ***
## Sex         -1.230e+00  8.215e-01  -1.497   0.1345    
## Age         -8.590e-02  1.389e+00  -0.062   0.9507    
## Age2        -5.001e-03  5.775e-02  -0.087   0.9310    
## AgeSex       1.385e-01  6.881e-02   2.012   0.0443 *  
## PC1          7.414e+00  1.234e+01   0.601   0.5480    
## PC2         -2.848e-01  1.497e+01  -0.019   0.9848    
## PC3          1.511e+00  9.654e+00   0.157   0.8756    
## ---
## Signif. codes:  0 '***' 0.001 '**' 0.01 '*' 0.05 '.' 0.1 ' ' 1
## 
## Residual standard error: 1.006 on 2113 degrees of freedom
## Multiple R-squared:  0.05444,    Adjusted R-squared:  0.05086 
## F-statistic: 15.21 on 8 and 2113 DF,  p-value: < 2.2e-16
```

```
effectsize::standardize_parameters(mediation_6)
```

```
# standardized indirect effect
RTV_M4 <- lm.beta(mediation_4)
RTV_M5 <- lm.beta(mediation_5)
RTV_ie <- (RTV_M4$standardized.coefficients["ADHD_PRS"])*(RTV_M5$standardized.coefficients["RTV"]) # times the standardized a path by the standardized b path
RTV_ie
```

```
##   ADHD_PRS 
## -0.0185943
```

```
# proportion mediated
RTV_M6 <- lm.beta(mediation_6) # standardized c (total) path
RTV_ie/(RTV_M6$standardized.coefficients["ADHD_PRS"])# divide the standardized indirect effect by the standardized total effect
```

```
##  ADHD_PRS 
## 0.1552446
```
